# Sex differences and multiregional variation in placental gene expression in late-onset gestational hypertension

**DOI:** 10.64898/2026.11.25.769234

**Authors:** Seema B. Plaisier, Melanie Smith, Aseel Abdulahad, Lora Nordstrom, Dean V. Coonrod, Claire T. Roberts, Melissa A. Wilson

## Abstract

Placental dysfunction has been associated with hypertensive disorders of pregnancy which are primarily characterized by high maternal blood pressure newly diagnosed after 20 weeks of gestation (gestational hypertension). Of these, early-onset preeclampsia (≤ 34 weeks) has dominated previous research because it is associated with more severe adverse outcomes for the mother and fetus, but late-onset preeclampsia with mild features or late-onset gestational hypertension (>34 weeks) is much more common and can be life-threatening for the mother during pregnancy and increase the risk of future health complications. As sex differences have been observed in the incidence of late-onset preeclampsia (more common when carrying a male fetus), we analyzed differential gene expression in placental tissue from full-term deliveries of ten late-onset gestationally hypertensive and ten normotensive pregnancies, each with nearly half carrying a fetus assigned female at birth (46,XX karyotype) and half assigned male at birth (46,XY karyotype). Every placenta was sampled at four evenly distributed locations, allowing analysis of intraplacental variation across the placental disc. Gene expression differences between hypertensive and normotensive pregnancies in female placentas were enriched for antigen receptor mediated signaling, response to angiotensin, cell junction disassembly, and ribosomal genes. In male placentas, gene expression differences between hypertensive and normotensive pregnancies were enriched for ribosomal genes as well as genes involved in mitochondrial function and DNA/RNA processing, immune system, and inflammatory functions. Comparisons with two previously published preeclampsia datasets identified overlapping placental gene expression signatures which included inflammatory and immune response genes in male placentas and cell adhesion and extracellular matrix organization genes in female placentas. Genes with increased intraplacental variation in gene expression in hypertensive pregnancies were enriched for muscle cell differentiation, vascular development, and response to mechanical stimulus, all related to the establishment of the feto-maternal interface. Previously reported placental marker genes of preeclampsia such as *LEP* and *FLT1* showed greater differential expression comparing hypertensive to normotensive pregnancies in male placentas than in female placentas and were among the highest in intraplacental variation. In summary, we show that differences in placental gene expression associated with late-onset gestational hypertension vary by fetal sex.

**Plain English summary:** The placenta is an organ that develops during pregnancy to supply the baby with oxygen and nutrients and remove waste as the baby’s organs mature. Because the placenta and baby are formed from the same fertilized egg, both have the same DNA, including sex chromosomes (XX for females or XY for males), and sex can affect the expression of certain genes in healthy and complicated pregnancies. We measured gene expression in 4 locations in 20 placentas, half from pregnancies where the mother developed high blood pressure in late pregnancy and half from mothers with normal blood pressure, where each group was half male-bearing and half female-bearing. We found that the genes that had different expression levels in placenta from high blood pressure pregnancies compared to normal blood pressure pregnancies were different in each sex. Placentas growing alongside female babies showed gene expression changes in high blood pressure pregnancies related to response to specific cell signals, while those alongside male babies showed changes related to energy production and metabolism, immune response, and cell adhesion. Previously reported marker genes of this condition were largely changed only in placentas from male-bearing pregnancies. Genes with more variability in expression across the placenta in high blood pressure pregnancies were involved in processes for establishing the interface between the placenta and maternal uterus, possibly indicating ways the placenta has struggled to function or responded to high blood pressure during pregnancy. In summary, sex is important for understanding how the placenta is affected in pregnancy complications.

**Highlights:**

- Differentially expressed genes in the placenta in pregnancies with late-onset gestational hypertension (preeclampsia without severe features) compared to normotensive pregnancies are distinct for each fetal sex
- Female placentas showed significantly decreased expression of ribosomal genes and increased expression of response to angiotensin, PDGFRβ signaling, Toll-like receptor signaling, and T cell receptor signaling
- Male placentas showed significantly decreased expression of ribosomal genes, mitochondrial processes, DNA replication, and amino acid metabolism and increased expression of complement pathway, humoral immune response, and cell adhesion-related genes
- Known preeclampsia placenta marker genes showed more differential expression in placenta from male-bearing pregnancies in three datasets for hypertensive disorders of pregnancy and were among the most variably expressed genes across the placental disc
- Genes with higher intraplacental variation in placentas from pregnancies with late-onset gestational hypertension were significantly enriched for vasculogenesis, axonogenesis, and muscle cell differentiation, all processes important to the establishment of a fetomaternal interface

## Background

The placenta is a vital transient organ formed to serve as the interface between a mother and a developing fetus during pregnancy. The placenta serves all the roles of the internal organs of the fetus while their organs grow and become functional, and protects the fetus from the maternal immune system. Trophoblast cells differentiate from the fertilized oocyte forming the outer layer of the blastocyst, which then invades into maternal tissue, forming the placenta as an interface that grows to adjust to the needs of the growing fetus throughout pregnancy. As the placenta and fetus both originate from the fertilized egg, the placenta and fetus share the same genome and thus share the same genetic sex. Sex differences in gene expression have been observed during development in the fetus and placenta as well as many other organs throughout the life span (1,2). While placental gene expression varies based on fetal sex, many placenta genomics studies do not specifically discuss whether the phenomena they are studying are different between sexes or investigate how these differences affect pregnancy complications (3,4). Sex differences in how the placenta develops, especially in the context of pregnancy complications, can have significant effects on the future development and progression of adult diseases for the fetus after birth, as well as cause poorer health outcomes for the mother during and after pregnancy (5).

Hypertensive disorders of pregnancy are common pregnancy complications that have been associated with placental dysfunction. Preeclampsia is the most common hypertensive disorder of pregnancy and is primarily characterized by arterial hypertension that presents during pregnancy after 20 weeks of gestation; additional features may include proteinuria, headaches, epigastric pain, visual disturbances, seizures, end-organ damage, and, if left untreated, preterm birth and maternal and/or fetal mortality (6,7). There are two types of preeclampsia: early-onset (≤ 34 weeks) and late-onset (>34 weeks), which are thought to be distinct clinical entities and have been linked to different types of placental dysfunction (8). Early-onset preeclampsia has been the subject of many placenta transcriptome studies because it often comes with a more severe clinical presentation, often including defective placentation due to poor invasion of extravillous cytotrophoblasts into the maternal uterine tissue which impairs remodeling in decidual and myometrial segments of the spiral arteries and leads to intrauterine fetal growth restriction (9). Late-onset preeclampsia typically does not present with defective spiral artery formation during placentation, but does present with impaired placental perfusion thought to be due to an imbalance of angiogenic factors such as *PlGF* and increased sFLT1 (10). Late-onset preeclampsia is seven times more common, may be linked to pre-existing maternal health conditions such as obesity, chronic hypertension, and diabetes, and is associated with poor birth outcomes, fetal morbidity or mortality, and later life cardiovascular disease, but is underrepresented in placenta research (11,12). Sex differences have been reported in both subtypes of preeclampsia, with female-bearing pregnancies being more often affected with early-onset while male-bearing pregnancies more often with late-onset preeclampsia (13–15). Retrospective analysis of gene expression in placental samples from cases from pregnancy cohorts collected in Australia showed statistically significant gene expression differences between late-onset preeclampsia and normotensive controls only in male-bearing pregnancies (16), demonstrating that placental gene expression differences in pregnancy complications are affected by fetal sex.

Studying gene expression in and across the placenta presents unique challenges. The placenta cannot be sampled when it causes potential harm to the mother or developing fetus; sample collection at delivery requires staff availability and coordination at unpredictable times and under unpredictable clinical circumstances (17). Placental and maternal tissue may be hard to separate (17). In the few studies that have sampled placentas at more than one location, the average interplacental variation in gene expression was larger than intraplacental variation, but substantial intraplacental variation was also observed (18,19). Variation in bulk tissue gene expression measurements can be driven by differences in cell type proportions across the placenta, normal variability in placental size and structure, and biochemical factors such as the relative levels of oxygen exposure (20,21). Additional studies are needed to understand which genes show high variability in expression across the placental disc and the impact of that variation on studying placenta in healthy and complicated pregnancies.

To address these important considerations, this study presents a sex-balanced, multiregional placenta gene expression dataset to characterize sex-specific gene expression changes in late-onset gestationally hypertensive pregnancies. Sex-balanced experimental design and sex-aware differential expression analysis allowed the identification of genes that are differentially expressed by sex in uncomplicated pregnancies and how expression is affected in gestationally hypertensive pregnancies within the same fetal sex. We found that genes differentially expressed between hypertensive and normotensive pregnancies were not well correlated between fetal sexes and previously published placental marker genes of preeclampsia were largely differential only in males. This study also highlights high intraplacental variation in genes that have been associated with placental function and hypertensive disorders of pregnancy, with genes that have higher intraplacental variation in hypertensive pregnancies showing enrichment for processes known to affect the establishment of the fetomaternal interface. Collectively, this study investigates how placental gene expression is altered in pregnancies with late-onset gestational hypertension in each fetal sex to determine sex-specific mechanisms underlying changes in placental biology in this pregnancy complication.

## Methods

### Valleywise Pregnancy Cohort

Working with the Valleywise Health Medical Center in Phoenix, Arizona (United States), we collected placenta samples from 20 pregnancies, half with normal blood pressure throughout gestation and half where mothers were diagnosed with late-onset gestational hypertension or preeclampsia without severe features after 34 weeks of gestation. Of the placenta samples from the 10 from normotensive pregnancies, 5 were male (assigned male at birth) and 5 were female (assigned female at birth). Of the 10 placenta samples from late-onset gestationally hypertensive pregnancies, 6 were male and 4 were female. Included mothers were between 20 to 34 years of age, low parity (nulliparous or 1), had White as their self-reported race, had Hispanic as their reported ethnicity, and delivered vaginally. Exclusion criteria included no history of chronic hypertension, no other pregnancy complications such as gestational diabetes, no substance abuse during pregnancy, and no active infections requiring antibiotic treatments except for group B strep.

### Placenta tissue sampling

All placentas were collected at full-term (≥ 37 weeks) delivery. Placental villous samples were taken midway between the chorionic and basal plates from the periphery of the lobules avoiding maternal tissue. Placentas were oriented maternal side up and the basal plate was removed to avoid sampling the maternal tissue. Four tissue samples, sampled equidistant in quadrants across the placenta, weighing roughly 50 mg each. Half of each sample was placed in a cryovial, snap frozen in liquid nitrogen, and stored at −80 °C and half of each sample was placed in RNAlater then stored at -80 °C. Chosen sampling sites were free of visible infarction, calcification, hematoma, and tears. All placenta samples were collected immediately following live birth via vaginal delivery.

### Whole exome sequencing

1 µg of genomic DNA is sheared to a mean fragment length of about 140 bp using focused acoustic energy (Covaris E210). Magnetic AMPure XP beads (Beckman Coulter) were used to purify the sheared DNA samples and remained with the sample throughout library construction. Samples yielding at least 1 μg of amplified DNA were used for capture. Sample concentrations were normalized to 2nM and loaded onto Illumina NovaSeq X Plus flow cells and sequenced using 101 bp paired-end sequencing reads according to Illumina protocols. Signal intensities were converted to individual base calls during a run using the system’s Real Time Analysis (RTA) software. Base calls were transferred to a high performance computing cluster and samples were de-multiplexed and aligned to the human genome using Illumina’s CASAVA 1.8.2 software suite.

### RNA sequencing

Using the Kapa RNA HyperPrep Kit with RiboErase (KR1351), ribosomal RNA was depleted starting from 25-1000ng of total RNA. Indexed libraries that met appropriate cut-offs for both quantity and quality were quantified by qRT-PCR using a commercially available kit (KAPA Biosystems) and insert size distribution determined with the LabChip GX or Agilent Bioanalyzer. Samples with a yield of ≥0.5 ng/ul were used for sequencing. Sample concentrations were normalized to 1.2 nM and loaded onto an Illumina NovaSeq X flow cell. Samples were sequenced using 100bp paired-end sequencing on an Illumina NovaSeq X according to Illumina protocols. Samples were sequenced to 50 million (M) 2 × 100 bp paired-end reads. Base calling, de-multiplexing, and alignment were done similarly to whole exome sequencing.

### Sequencing data processing

Samples were checked for quality using FastQC (v 0.11.8) (22) and aggregated using MultiQC (v 0.9) (23). For preprocessing of the RNAseq data, we trimmed adapters using bbduk as part of bbmap (v 38.22) (24) with the following parameters: qtrim=rl trimq=30 minlen=75 maq=20. Post trimming quality was again checked using FastQC (v 0.11.8) (22) and MultiQC (v 0.9) (23). Post trimming samples had an average of 35.18 M and median of 35 M reads. After verifying that samples from male placentas had higher coverage of the Y chromosome and samples from female samples did not, all sequencing samples were aligned to Gencode GRCh38.p12 human reference genome informed on the sex chromosome complement of the sample (25,26). Samples with no evidence of a Y chromosome were aligned to a reference genome with the entire Y chromosome masked with Ns to avoid mis-mapping of homologous X–Y sequence reads (25). Samples with evidence of a Y chromosome were aligned to a reference genome with the Y chromosome pseudoautosomal regions (PARs) masked out as those regions are replicated 100% on the X chromosome PARs in GRCh38.p12. For exome sequencing, reads were aligned using bwa (v 0.7.17) and variants were called using gatk (v 4.2.6.1). For RNA sequencing, reads were aligned using HISAT2 (v 2.1.0) (27) and gene expression was quantified using SubRead featureCounts (v 1.5.2) (28). We followed the XY_RNAseq protocol (25) to utilize a Ymasked (for XX samples) and YPARs masked (for XY samples) reference genome and HISAT -x index function to create two sex chromosome complement reference indexes used for alignment. HISAT2 alignment was performed with the following parameters, --dta for downstream transcriptome assembly, --rna-strandness RF to indicate the sequences are reverse forward, --phred 33 encoding, and pair-end alignment. RNAseq alignment files were then sorted, read groups were added, duplicates were marked, and files were indexed using bamtools 2.5.1 (29) and Picard 2.9.2 (30). FeatureCounts was employed using --primary to only use primary alignments and -p 2 to specify the minimum number of consensus reads from the same pair, suggested for paired-end read data (28). FeatureCounts uses the gene annotation file to infer exon–exon junctions from connecting each pair of neighboring exons from the same gene. FeatureCounts was run for each RNAseq sample, at the gene level using -g gene_name. There are 57,133 genes in the Gencode GRCh38.p12 human reference genome used in this analysis.

### Genetic ancestry prediction

Whole exome sequencing was used to predict genetic ancestry using peddy (31). All samples were predicted to be from the same genetic ancestry group, AMR (Admixed American), so we did not include this as a covariate in our differential expression analysis.

### Chromosomal copy number variation

Whole exome sequencing was used to detect copy number variation using CNVkit (v 0.9.11). As our placenta samples were not paired, a pooled reference was created using the ‘batch’ tool with aligned whole exome sequencing reads from all 20 placenta samples, capture target files for the KAPA HyperExome V2 platform downloaded from Roche sequencing (https://sequencing.roche.com/content/dam/diagnostics_microsites/sequencing/master-blueprint/en/products-solutions/sample-prep/dna-reagents/target-enrichment/hybridization/hyperexome-probes/v2/KAPA-HyperExomeV2-design-files_hg38.zip), the GRCh38.p12 human genome reference filtered for only assembled chromosome sequences, and an accessible regions file created using the ‘access’ tool with the reference genome sequence file. This pooled reference was compared to all samples using the ‘batch’ tool and the output was visualized using the ‘scatter’ tool. Seeing no evidence of large scale chromosomal variation in any sample, chromosome log2 ratios were summarized by taking a weighted mean of log2 of all segments in each chromosome (‘weighted.mean’ function in R v 4.5.0).

### Imputed cell fractions

To input cell fraction of placental/maternal cell types, cell type signatures were created using single cell and single nucleus RNA sequencing results published by Admati *et al.* (32). These data sets were chosen because they included samples taken from term deliveries, as our samples were. The Admati *et al.* data set was filtered for only late pregnancy and vaginal delivery samples which matches our data set and single cell RNA sequencing data was available for download. The Admati *et al.* data set included placentas from both preeclamptic pregnancies and normotensive uncomplicated pregnancies, cells clustered according to cell type in both cases and were aggregated to get a single cell reference matrix for deconvolution. Counts were loaded to a Seurat object and filtered with the following criteria: minimum cell threshold of at least 3, at least 200 RNA features, no greater than 2500 RNA features, and percent mitochondrial less than 5. Data was normalized (‘NormalizeData’, Seurat, v 5.3.0), scaled (‘ScaleData’), and the top 5 principle components were used to visualize the data using a UMAP (‘FindNeighbors’ dims = 1:20, ‘FindClusters’ resolution = 0.5, ‘RunUMAP’ dims = 1:20) to make sure cells showed clustering by cell type. Data was loaded into a SingleCellExperiment object and aggregated to be passed into CIBERSORTx (‘aggregateData’, SingleCellExperiment, v 1.30.1). Genes were filtered for those that were expressed in at least 10% of the aggregated cell types. Cell type signature matrices were created using CIBERSORTx ‘Create Signature Matrix’ tool on the web application (https://cibersortx.stanford.edu/about.php) (33). These signature matrices were passed into the ‘Impute Cell Fractions’ module of CIBERSORTx with the normalized log_2_-transformed counts per million of all genes across all of our Valleywise samples as the ‘Mixture File’. Permutation analysis was used to assess significance with 1000 permutations. All deconvolutions had p-values of correlation below 0.1 indicating that the single cell reference profiles were confidently identified in our bulk gene expression profiles. The statistical significance of differences between cell type proportions between males and females and between hypertensive and normotensive pregnancies were assessed using linear regression which accounted for multiple sampling from the same placenta using the ‘lmer’ function to fit a model with two groups (lme4, v 1.1-38) and the emmeans function to assess significance (emmeans, v 2.0.0).

### Published Preeclampsia Data Sets

Gene expression profiles were compared to published placenta RNA sequencing datasets: Horii *et al.* 2023 (34) and Smith *et al.* 2025 (16). The Smith dataset included sex-stratified gene expression profiles created using similar techniques with their publication. We reanalyzed the Horii dataset after sex-stratification using the limma-voom method so that analysis stayed consistent across data sets. Counts were downloaded from Gene Expression Omnibus (GSE186257), normalized with TMM, converted to log_2_ CPM, and analyzed for differential expression using limma-voom for consistency with Valleywise placenta gene expression analysis.

## Statistical Analysis

### Quantifying differences in clinical features in enrolled study participants

We examined sex differences for clinical information for placentas for maternal age at delivery, paternal age at delivery, pre-pregnancy maternal body mass index (BMI), paternal BMI, gravidity, parity, and gestational age at delivery. Continuous variables were tested for sex differences using a Welch’s two-sample t-test. Categorical variables were tested for sex differences for categorical variables using a Pearson’s chi-squared test. The threshold for statistical significance was being under a p-value of 0.05.

### Differential gene expression analysis

Sex-differential expression analysis between male (XY) and female (XX) placentas was performed using the limma/voom pipeline (35,35). Quantified read counts from each sample were generated using SubRead featureCounts and combined into a count matrix, with each row representing a unique gene ID and each column representing the gene counts for each unique sample. The counts matrix was loaded into a DGEList object using the limma package (v 3.66.0). Genes with low expression in the groups being compared were filtered with the ‘filterByExpr’ function in the edgeR package (v 4.8.2). Normalization factors were calculated using the calcNormFactors function in the edgeR package (36). Then, we ran Trimmed Means Method (TMM) to normalize for library size variation between samples (37). Counts were then transformed to log2(CPM + 0.25/L), where CPM is counts per million, L is library size, and 0.25 is a prior count to avoid taking the log of zero (38). Correction for the sequencing batch was performed by including the batch as a covariate during differential expression as well as a placenta identifier to account for multiregional sampling of each placenta (∼comparison_group + batch + (1 | placenta_id)). Voom transformation was conducted using the ‘voomWithDreamWeights’ function, the model was fitted using the ‘dream’ function, and finally an empirical Bayes approach was applied to determine differentially expressed genes using the ‘eBayes’ function (variancePartition v 1.40.1).

### Pathway enrichment analysis

Differentially expressed gene sets defined using specific statistical thresholds were analyzed for enrichment of Gene Ontology (GO) Biological Processes using clusterProfiler (v 4.18.4) (39–41).

### Gene Set Enrichment Analysis (GSEA)

Gene set enrichment analysis was implemented in a threshold-free manner using fgsea (v1.36.2). Genes were ranked based on differential expression results: the modified t-statistic (‘t’) or log-transformed fold change (‘logFC’) as output by the topTable function in limma (v 3.66.0). Enrichment was measured for gene sets from three collections downloaded using the msigdbr package (v 26.1.0): hallmark gene sets (collection = “H”), Gene Ontology biological processes (collection = “c5”, subcollection = “GO:BP”), and KEGG pathways (collection = “C2”, subcollection = “CP:KEGG_LEGACY”). Enrichment results for the selected gene set collection and ranked gene list were attained using the ‘fgsea’ function (minSize = 15, maxSize = 500). Enrichment results were filtered for by thresholding the adjusted p-value (FDR-controlled p-value). Network views of the significantly enriched gene sets were created using ggraph (v 2.2.2) and igraph (2.3.3).

### Rank-rank hypergeometric overlap (RRHO)

RRHO analysis was implemented using the RRHO package (v 1.50.0). All genes were ranked by log-transformed fold change (‘logFC’) for each placenta dataset, comparing placenta from hypertensive pregnancies to those from normotensive pregnancies separately for male and female placentas. For every pair of ranked gene lists being compared, genes were filtered for those present in both gene lists being compared. RRHO analysis was performed using the ‘RRHO’ function (alternative = “enrichment”, stepsize = 100, plots = TRUE, BY = TRUE). Output included the RRHO map showing areas of significant overlap and gene lists of the significant overlap in the UpUp and DownDown regions. These gene lists were analyzed for functional enrichment using clusterProfiler (see above).

### Intraplacental variation

For each gene, the mean and variance of normalized, batch corrected gene expression (log CPM) across all 4 samples from each placenta were calculated (using ‘rowMeans’ and ‘var’ function in base R v 4.5.0). Then, a mean-variance trend was fit using the ‘loess’ function (log10(gene variance) -gene mean, span = 0.5). The residual variability was attained using the ‘residuals’ function on that fit (stats package from R v 4.5.0).

## Results

### Sex-balanced, multiregional placenta sampling to investigate gene expression differences across sexes and late-onset gestational hypertension

To assess sex differences in placental gene expression and how those sex differences change in pregnancies with late-onset gestational hypertension, a sex-balanced experimental design was implemented. Of the 20 placentas sampled during our study, half were from normotensive, uncomplicated pregnancies and half were from pregnancies with late-onset gestational hypertension, each with nearly half carrying a male fetus and half carrying a female fetus with sex assigned at birth (Figure 1, Supplemental Table 1). All placenta samples were collected at full-term vaginal delivery and carefully dissected to collect villous tissue in areas that did not show signs of calcification or tissue damage. Mothers who enrolled in the study were between 20 to 34 years of age, and of the same self-reported race and ethnicity (White and Hispanic, respectively). Most mothers (18 of 20) had an elevated body mass index (> 25). Enrollment was limited to mothers that did not have a history of elevated blood pressure outside of pregnancy. There was no statistically significant difference in features between pregnancies with different gestational hypertension status (Supplemental Table 2). Maternal and paternal ages were about 4 years younger in male-bearing pregnancies than female-bearing pregnancies; there was no statistically significant difference in gravidity, parity, maternal BMI, paternal BMI, or gestational age at delivery by fetal sex. Genetic ancestry was inferred and all samples were observed to be of Admixed American ancestry (AMR) (Supplemental Table 3). Chromosome copy number estimates from exome sequencing data showed no evidence of large scale differences in copy number for any chromosome in any placenta (Supplemental Table 4).

**Figure 1.**
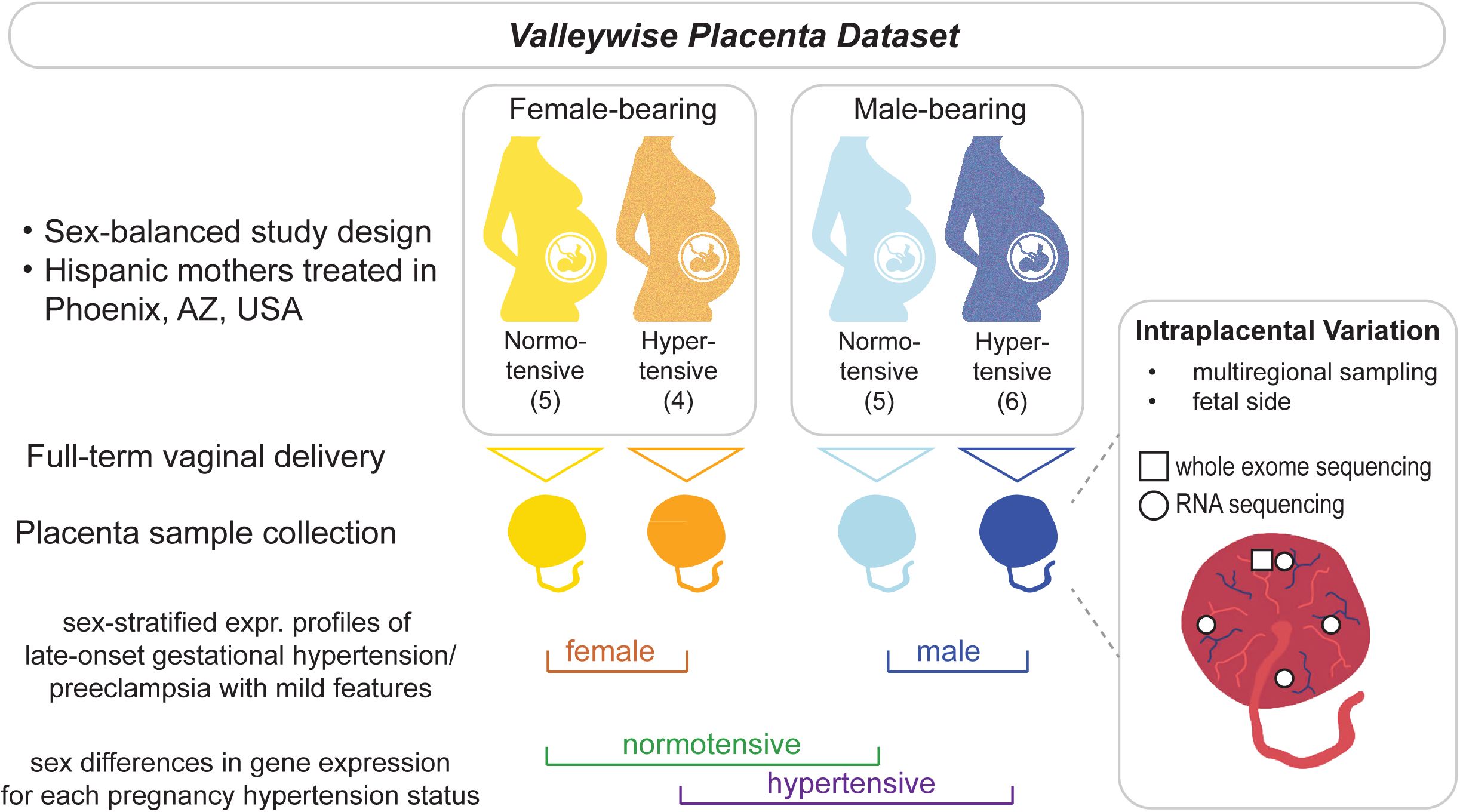
Experimental and analytical design. The Valleywise placenta dataset employed a sex-balanced study design, wherein placentas were carefully dissected on the fetal side after full-term vaginal delivery from half female-bearing and half male-bearing pregnancies. For each fetal sex, half of the pregnancies were diagnosed with gestational hypertension and/or late-onset preeclampsia. Each placenta was sampled once for exome sequencing and 4 times for gene expression using RNA sequencing. Profiles of gene expression were analyzed by doing pairwise comparisons between 4 groups accounting for fetal sex and hypertension status and intraplacental variation in gene expression was calculated for every placenta sampled.

### Hypertension-Associated Placental Gene Expression Profiles Differ by Fetal Sex

We first asked whether sex differences in normotensive pregnancies were replicated under the hypertensive condition. We observed a positive correlation (R=0.6; p-value < 2 x 10^-16^) in sex differences between placentas from hypertensive and normotensive pregnancies (Figure 2A). Genes significantly differentially expressed between sexes in both hypertensive and normotensive pregnancies after multiple hypothesis correction (adj. p-value < 0.05) were almost all sex chromosome genes (Supplemental Table 5). We observed a total of 51 genes that show differential expression by sex, of which 32 exhibit sex differences in both hypertensive and normotensive pregnancies, 16 in normotensive pregnancies only, and 3 in hypertensive pregnancies only (Supplemental Fig 1A,C). Genes that had a high male to female expression ratio were nearly all chromosome Y genes, which was expected as female placentas do not have a Y chromosome and as such the gene expression data was analyzed using sex-aware alignment methods. Similarly, genes with the most negative male to female expression ratio were XIST and TSIX, which are involved in X chromosome inactivation, and thus expected to be highly expressed only in female placenta tissues. Excluding genes on the X and Y chromosomes, the correlation is reduced but still statistically significant (correlation coefficient = 0.24, p-value < 2 x 10^-16^) (Figure 2A). While sex differences in gene expression were mostly in sex chromosome genes, there were 9 autosomal genes that were differentially expressed between sexes in either normotensive or hypertensive pregnancies (Figure 2B, Supplemental Table 5). Of particular note is *PDGFA* which is known to regulate cell proliferation and blood vessel growth. Growth factors such as this have been shown to be upregulated in trophectoderm cells during implantation and in cytotrophoblasts at the base of growing placental villi (42,43). *PDGFA* was more highly expressed in male placentas than female placentas from normotensive pregnancies. *PDGFA* did not show differential expression by sex in placentas from hypertensive pregnancies, where *PDGFA* was equally lowly expressed in both sexes, more similar to the level observed in normotensive female placentas (Figure 2). This may indicate a different balance of angiogenic factors between sexes in normotensive pregnancy that is altered in hypertensive pregnancies. Other genes with sex differences in normotensive but not hypertensive pregnancies included *DDX55*, which had higher expression in males than females from normotensive pregnancies, and *PCP4L1*, *DGCR8*, *TAOK2*, *PRRG2*, and *HMG20B*, which had higher expression in females than males from normotensive pregnancies. Genetic variations or altered methylation of several of these genes in the placenta have been associated with pregnancy complications, but no clear mechanism has been suggested (44–48). In hypertensive pregnancies, *IQSEC3* showed higher expression in female placentas than in male placentas but was not differentially expressed by sex in placentas from normotensive pregnancies. *IQSEC3* is an ARF guanine-nucleotide exchange factor that has been linked to placental abruption in genome wide association studies (47). *EIF1AXP1* is a translation initiation factor that we observe has higher expression in female than male placentas in both hypertensive and normotensive pregnancies and has been previously shown to have significantly higher expression in female placentas from uncomplicated pregnancies (1). Interestingly, *EIF1AXP1* is found on chromosome 1 but it is a processed pseudogene from chromosome X and significantly higher expression in females has also been observed in human aortic smooth muscle cells (49). These results suggest that placenta expression in normotensive pregnancy varies based on sex. Future studies should consider the sex of the samples when studying variation between normotensive and disease conditions.

**Figure 2.**
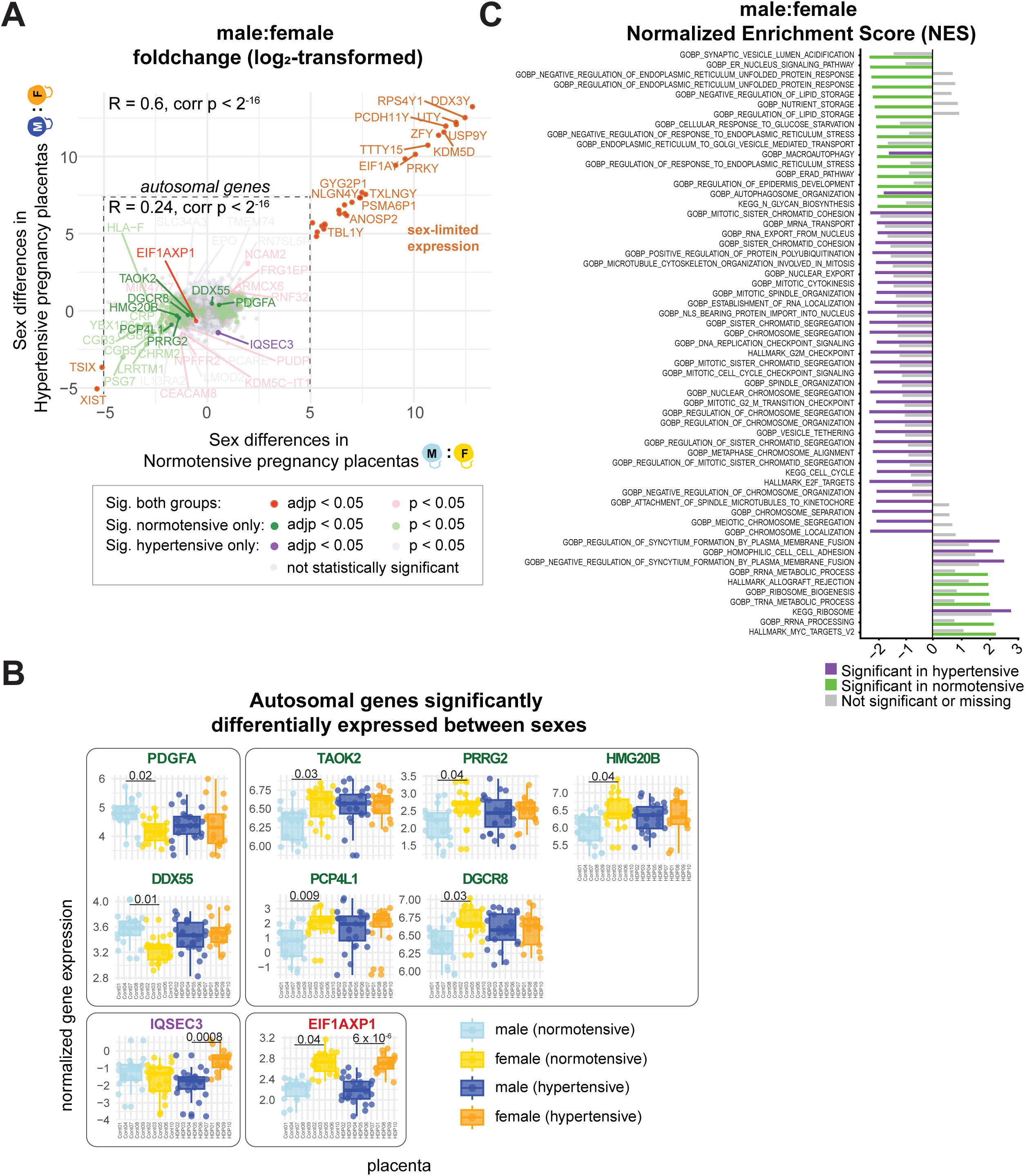
Differential gene expression in placenta between sexes in each pregnancy hypertension group. (**A**) Scatter plot showing the log_2_ transformed fold change in male placentas versus female placentas in normotensive pregnancies (x-axis) compared to hypertensive pregnancies (y-axis). (**B**) Violin plots showing normalized expression levels for autosomal genes that are significantly differentially expressed (adjusted p-value < 0.05). Boxes indicate a similar pattern of expression between genes. (**C**) Significantly enriched gene sets (adjusted p-value < 0.05, normalized enrichment score >= 1.9) for differential expression between male placentas versus female placentas in hypertensive pregnancies (purple) and normotensive pregnancies (green). Positive NES score indicates higher expression of genes in the enriched gene set in male placentas; negative NES score indicates higher expression in female placentas.

As our study focused on placental gene expression at full-term delivery, it represented a more common, but often less severe, hypertensive disorder of pregnancy. We therefore anticipated that changes in gene expression would be more subtle and that coordinated, smaller scale changes in multiple genes that modulate the same cellular processes may underlie differences in placental function in this pregnancy complication in each fetal sex. We used gene set enrichment analysis (GSEA) to find significant concordant expression of genes involved in various biological processes and signaling pathways. Genes were ranked by moderated t-statistic in each of our four differential expression profiles, so as to represent the difference in average expression attenuated for variation in the two groups being compared. GSEA was run using three collections of genes from the Molecular Signatures Database (MSigDb): Hallmark gene sets, Gene Ontology Biological Process gene sets, and KEGG pathway gene sets (Figure 2C and 3C,D, Supplemental Tables 5 and 6). Enrichment results indicate that in normotensive pregnancies, female placentas have higher expression of genes involved in lipid and nutrient storage and endoplasmic reticulum stress, while males have higher expression of genes involved in RNA processing. Hofbauer immune cells in female placentas at term have been shown to have more efferocytosis (biological process by which dying or apoptotic cells are recognized, engulfed, and digested by phagocytic cells) powered by lipid metabolism and showed endoplasmic reticulum stress response, while males showed more glycolytic metabolism (50). In hypertensive pregnancies, genes related to mitosis, cell cycle, and chromosome segregation had increased expression in females, while genes related to syncytium formation and cell-cell adhesion have increased expression in males (Figure 2C). In both hypertensive and normotensive pregnancies, autophagosome genes were significantly enriched in genes with higher expression in females. Previous studies in a mouse model with an essential autophagy gene deleted exhibited different sex-specific responses to obesity including decreased expression of mitochondrial genes, consistent with our results in male placentas from hypertensive male pregnancies (51). These results point to sex-specificity in gene expression that may underlie sex differences in placenta biology in pregnancies with and without late-onset hypertension.

**Figure 3.**
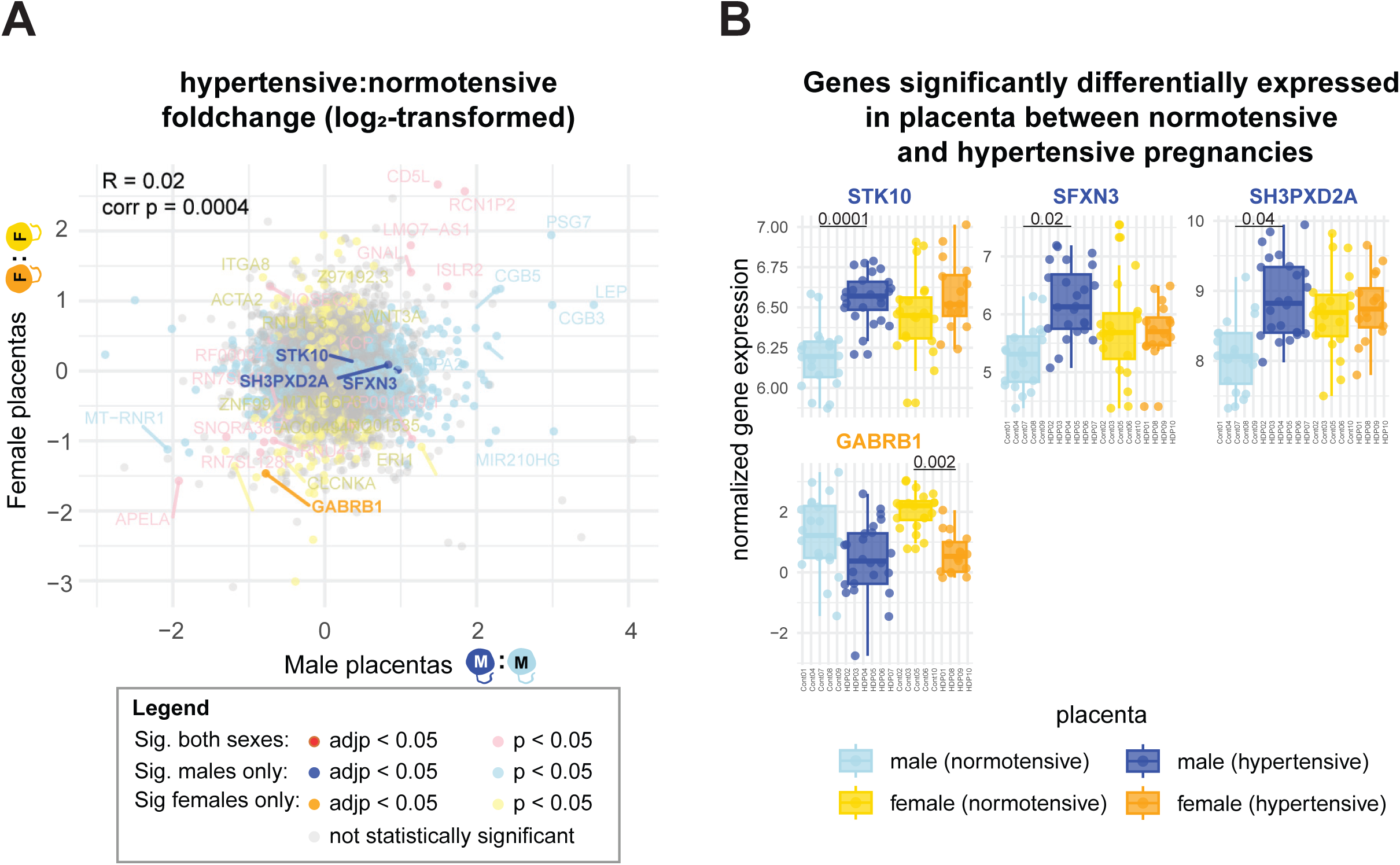
Differential gene expression in placenta between pregnancy hypertension groups in each sex. (**A**) Scatter plot showing the log_2_ transformed fold change in placentas from hypertensive pregnancies to normotensive pregnancies in males (x-axis) compared to females (y-axis). (**B**) Violin plots showing normalized expression levels for genes that are significantly differentially expressed (adjusted p-value < 0.05).

Having compared male to female placentas within pregnancies with the same gestational hypertension status, we assessed differentially expressed genes in placentas from hypertensive compared to normotensive pregnancies within the same sex. Sex-stratification yielded more significantly differentially expressed genes than comparing all placentas from hypertensive pregnancies to all from normotensive pregnancies, particularly in males where the number of differentially expressed genes was nearly three times higher and included many placental markers of preeclamptic pregnancies (Supplemental Figure 1 G). The correlation between sex-stratified gene expression profiles of normotensive versus hypertensive pregnancies is statistically significant but very close to 0 (correlation coefficient = 0.02, p-value = 0.0004), indicating that the profiles of placental gene expression differences between hypertensive and normotensive pregnancies were different within each fetal sex. Using multiple hypothesis testing corrected thresholds for differential expression between hypertensive and normotensive pregnancies (adjusted p-value < 0.05), we observed three genes that were statistically significant in male placentas (*STK10*, *SFXN3*, and *SH3PXD2A*) and one gene in female placentas (*GABRB1*) (Figure 3 A,B). All three genes differentially expressed in males specifically, *STK10*, *SFXN3*, and *SH3PXD2A*, show higher expression in male placenta from hypertensive pregnancies compared to those from normotensive pregnancies with no statistically significant difference in female placentas. *STK10* is a serine/threonine kinase involved in cell adhesion and migration; oocyte-derived DNA methylation has been shown to affect *STK10* expression and trophoblast cell adhesion in murine placenta (52). *SFXN3* is a mitochondrial L-serine transmembrane transporter. A previous study has shown that *SFXN3* and other genes associated with oxygen-sensing subtype of syncytiotrophoblasts including *FLT1*, *PAPPA2*, and many *PSG* hormones involved in angiogenesis were upregulated when specific chromatin remodeling and transcriptional activator genes were knocked out in syncytiotrophoblast organoids (53). *FLT1*, *PAPPA2*, and *PSG 1/2/3/4/56/7/8/9/11* all showed higher average expression in male placentas from hypertensive pregnancies compared to normotensive, many with nominal statistical significance (unadjusted p-value < 0.05) (Supplemental Table 6). *SH3PXD2A* encodes a scaffold protein that has been previously shown to be differentially expressed and differentially methylated in placenta tissues from preeclamptic pregnancies along with *LEP* (54). *LEP* is also upregulated specifically between hypertensive and normotensive pregnancies in males, reaching nominal significance in our study (log_2_ fold change = 3.54, nominal p-value = 0.02) (Figure 3A, Supplemental Table 6). In contrast to males, only one gene, *GABRB1,* shows significantly decreased expression in female placentas from hypertensive pregnancies compared to normotensive pregnancies (adjusted p-value = 0.002). Although *GABRB1* shows a decrease in average expression in male placentas from hypertensive pregnancies compared to from normotensive, it exhibits increased intraplacental variation in all male placentas, which reduced the statistical significance of this separation (Figure 3B, Supplemental Table 6). *GABRB1* encodes a subunit of the GABA-A receptor which is involved in nervous system development and was shown to have significantly higher expression in male placenta tissues due to hypermethylation in female placenta tissues in the first trimester of pregnancy (55). In this study of placenta collected at term, there is no significant difference in expression between *GABRB1* between male and female placentas within normotensive pregnancies or within hypertensive pregnancies, but there is a decrease in expression in hypertensive pregnancies compared to normotensive within each fetal sex. *GABRB1* has been previously shown to be downregulated in neonates with adverse health outcomes where a sex-specific placental inflammatory response was observed (56). These genes may be involved in sex-specific differences in how the placenta causes or responds to hypertensive disorders of pregnancy.

Gene set enrichment analysis was again used to find concordant gene expression changes that might indicate genes associated with specific cellular functions that were different in placenta from hypertensive pregnancies versus normotensive pregnancies in each fetal sex. Significantly enriched gene sets (FDR-corrected enrichment p-value < 0.05) were filtered for those that had at least 60% of the gene set in the leading edge (genes that are contributing to the enrichment signal) (Figure 4 A,B). In both female and male placentas, genes with lower expression in those from hypertensive pregnancies were enriched for ribosomal genes, which are involved in the negative regulation of syncytium formation by plasma membrane fusion (Figure 4 A, B). This enrichment is driven by the reduced expression of many ribosomal protein genes (small and large). Syncytial sprouts, protrusions of syncytiotrophoblasts enlarging to grow the surface of exchange in a growing placenta, tend to have higher amounts of endoplasmic reticulum and ribosomes (57). Reduced expression of ribosomal proteins may represent an impaired protein translation that is necessary for the development of the syncytium and supporting the development of the fetus. Reduction of ribosomal genes has also been observed in placenta for which the fetus has intrauterine growth restriction (58).

**Figure 4.**
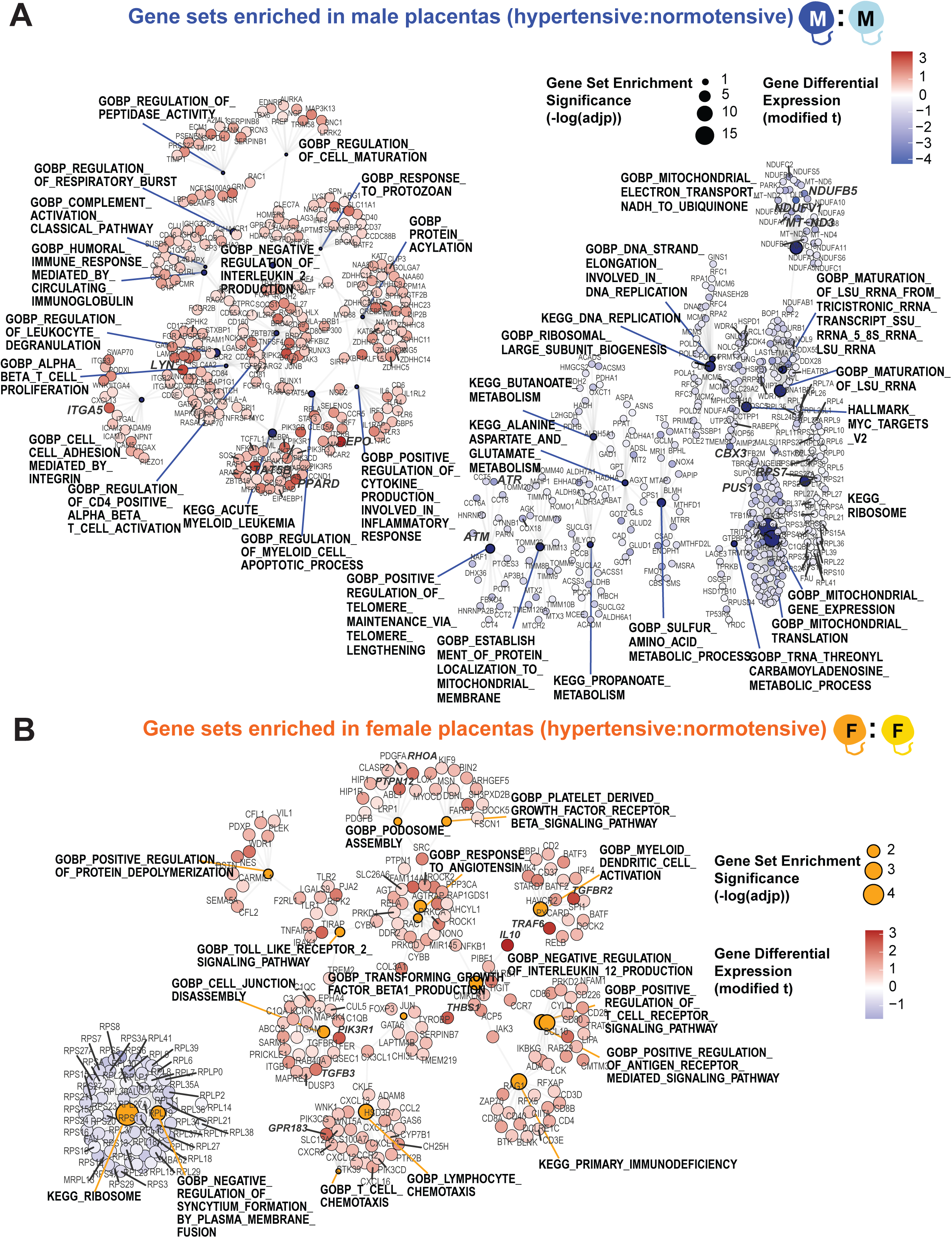
Significantly enriched gene sets for each sex based on gene expression differences in placenta from hypertensive versus normotensive pregnancies. Significantly enriched gene sets (FDR-controlled p-value < 0.05) that had at least 60% of the gene set as leading edge genes are shown for (**A**) male placentas and (**B**) female placentas. Nodes for significantly enriched gene sets are larger based on increased statistical significance (-log(FDR-controlled p-value)). Leading edge genes for each significant gene set are shown as nodes and nodes are colored according to the differential expression between placentas from hypertensive versus normotensive pregnancies (modified t-statistic from differential gene expression analysis); red indicates higher expression in placentas from hypertensive pregnancy and blue indicates lower expression in hypertensive.

We also found unique biological processes enriched between placentas from hypertensive versus normotensive pregnancies for each sex. In male placentas, genes with lower expression in placentas from hypertensive pregnancies were enriched for genes involved in mitochondrial transcription and translation, ribosomal genes, and amino acid metabolism (Figure 4A). Mitochondrial transcription and translation enrichment was due to lower expression of mitochondrial ribosomal protein genes (*MRP* genes) in male placentas from hypertensive pregnancies; these are genes that are on the nuclear genome (not the mitochondrial genome) and encode the protein subunits of mitochondrial ribosomes. Previous studies have reported lower expression of mitochondrial genes in placenta from preeclamptic pregnancies (67–72), but there was no indication whether this was sex-specific. In mice, male placentas from obese dams have reduced expression of mitochondrial genes (73,74). In our pregnancy cohort of mothers with higher body mass index, gestational hypertension is associated with lower expression of genes that are involved in mitochondrial function in male placentas. This includes genes involved in mitochondrial transcription and translation and transport to mitochondria. Gene sets involved in metabolism of amino acids and ion transport also have lower expression in male placenta from hypertensive pregnancies compared to normotensive, including many *NDUF* (NADH:ubiquinone oxidoreductase) genes. Male placentas also show downregulation of genes involved in DNA replication, which, along with the energy production from mitochondria, is needed for cell proliferation. Male placentas from hypertensive pregnancies show higher expression of genes involved in immune response, including genes that regulate interleukin production, T cell proliferation, and T cell activation. These differences in gene expression between placentas from hypertensive pregnancies compared to normotensive pregnancies may highlight mechanisms that are altered specifically in male placentas.

Genes increased in hypertensive pregnancies in females were enriched for genes involved in antigen receptor mediated signaling, immunodeficiency, Toll-like receptor signaling, cell junction disassembly, and response to angiotensin (Figure 4B). Antigen receptor mediated signaling, immunodeficiency, and Toll-like receptor signaling reflect immune modulation in the placenta. Published studies show that activation of Toll-like receptors systemically in pregnant rats induced preeclampsia-like symptoms (59). *TLR4* has been shown to be upregulated in preeclamptic placentas in trophoblast cells and Hofbauer cells (60,61). There is a heightened innate immune responsiveness in the third trimester of pregnancy, but this increase may be more dramatic in female placentas and may contribute to preeclamptic phenotypes in female-bearing pregnancies. Alterations in cell junction genes have been implicated in the formation of the syncytium while integrins have been reported to be lower in the placenta from preeclamptic pregnancies as a result of altered proliferative, invasive, and adhesive capacity of trophoblasts (62,63). The renin-angiotensin system has an important role in maintaining maternal blood flow into the placenta and with blood pressure regulation (64). Alterations in the components of this system in female placentas are thought to contribute to hypertension during pregnancy. Polymorphisms in the genes contributing to this enrichment signal including *AGT* (angiotensinogen) are linked to increased risk for preeclampsia and are linked to oxidative stress and with placental hypoxia causes an increase in reactive oxygen species which activate the NF-kB/RelA inflammatory pathway (64–66). Female placentas also showed higher expression of genes involved in immune response in hypertensive pregnancies but male placentas showed concurrent high expression of genes involved in inflammatory response, complement cascade, and defense response. This suggests that while both female and male placentas may have an altered immune state in hypertensive pregnancies, male placentas also show signs of acute stress with mitochondrial and metabolic dysfunction.

We next investigated whether these gene expression differences were due to differences in cell type proportions across the placenta samples. Single cell and single nucleus RNA sequencing data sets from placenta tissues sampled from full-term vaginal deliveries were used to create cell type reference matrices (32) and cell type proportions were inferred using single cell deconvolution (33) (Supplemental Figure 2, Supplemental Table 10). Single nucleus RNA sequencing was used to provide a better estimate of the proportion of syncytiotrophoblasts since the syncytium is generally removed during single cell RNA sequencing library preparations due to its multinucleated structure. The other placental cell types were represented with higher depth of coverage in single cell RNA sequencing (32). Cell types inferred to have the highest proportion in all placentas were trophoblasts (including syncytiotrophoblasts and villous cytotrophoblasts) and stromal cells. When samples were divided in 4 groups based on fetal sex and hypertension status, no pairwise comparison yielded significant differences in any cell type identified (Supplemental Figure 2A, Supplemental Table 10). When placenta from males and females were combined, there were cell types that show slight but significant differences between hypertensive and normotensive pregnancies. Using inference based on single nucleus RNA sequencing, the proportion of Hofbauer cells was inferred to be slightly lower in hypertensive pregnancies (p-value = 0.02), although the highest inferred proportion of this cell type in any placenta was only 1.1% (Supplemental Figure 2A, Supplemental Table 10). Using inference based on single cell RNA sequencing, all placentas from hypertensive pregnancies have a slightly higher inferred proportion of stromal cells that express STEAP4 (1.5% compared to 1.2% in normotensive pregnancies) (Supplemental Figure 2B, Supplemental Table 9). If placenta from hypertensive and normotensive pregnancies were combined, female placentas have a slightly higher inferred proportion of mature syncytiotrophoblasts (TB_SCT-PSG, p-value = 0.048). However, syncytiotrophoblast proportion inferred using single nucleus RNA sequencing data showed no significant difference between the sexes, so measurement is unreliable due to technical artifacts stemming from most cells in multinucleated syncytium being lost in single cell RNA sequencing assay protocols. Collectively, there was no substantial difference in inferred cell type proportion between placentas of different sex or pregnancy hypertension status in our dataset.

Given consistent inferred cell type proportions in our placentas grouped by fetal sex and pregnancy hypertension status, we used our dataset to more broadly investigate gene expression patterns between the four groups. Normalized expression of genes that were nominally significantly differentially expressed (p-value < 0.01) in at least one of our 4 differential expression profiles were hierarchically clustered by their expression pattern in all placentas in our dataset (male placenta from normotensive pregnancies, male placenta from hypertensive pregnancies, female placenta from normotensive pregnancies, female placenta from hypertensive pregnancies) (Supplemental Figure 3). The top six branches were used to represent the major patterns of gene expression that were differentially expressed between at least 2 of our 4 sample groups (Figure 5). Taking the mean expression of all genes in each cluster and of all the samples in each placenta, patterns of gene expression that span sex and hypertension status were observed (Figure 5). Cluster 1 genes had higher expression in male placentas from both hypertensive and normotensive pregnancies; this cluster included many chromosome Y genes. Cluster 1 genes were significantly enriched (adjusted p-value < 0.05) for oxidative stress related genes, including *HIF1A*, *IL10*, *MMP2*, and *FGF2*. This may indicate that male placentas were under greater oxidative stress than female placentas with or without hypertension during pregnancy. Clusters 2, 4, and 5 included genes where only males showed differential expression between placentas from normotensive versus hypertensive pregnancies. Cluster 2 genes showed higher expression in males from normotensive pregnancies than any other sample group. This cluster of genes was enriched for ribosome biogenesis, generation of precursor metabolites and energy, RNA metabolism and processing, and mitochondrial gene expression; this was consistent with previous studies that show that expression of these genes in males support increased growth during gestation (75). The expression of these genes was lower in male placentas from hypertensive pregnancies compared to normotensive pregnancies but their expression in females was similarly low in both hypertensive and normotensive pregnancies. These genes are interesting because they showed similarly low expression levels in male and female placentas from hypertensive pregnancies, but this expression level was different from placentas from normotensive pregnancies only in the case of males. Cluster 3 showed genes that were differentially expressed in female placentas from hypertensive versus normotensive pregnancies, but had intermediate expression in male placentas that was similar in hypertensive and normotensive pregnancies. This cluster did not show significant enrichment of any gene functions but did include the most differentially expressed genes in females placentas from hypertensive versus normotensive pregnancies, with two genes that had to do with the GABA receptor, *GABRB1* and *GABBR2*. GABA receptor expression in the placenta has been associated with placentation, brain development, hormone regulation, oxidative stress, and apoptosis of trophoblastic cells (74–77); these genes might contribute to the female placenta-specific effects of gestational hypertension. Clusters 4 and 5 comprise genes that had higher expression in hypertensive pregnancies in male placentas compared to normotensive, but showed no difference in expression in female placentas from hypertensive versus normotensive pregnancies. Cluster 4 genes had higher expression in the female placentas overall, whereas cluster 5 genes had expression in females at an intermediate state between male placentas from hypertensive and normotensive pregnancies. Cluster 5 genes were significantly enriched for protein catabolic process with *ATP13A2*, *AP5Z1*, and *MGAT3* in the leading edge. This cluster also included widely reported placenta preeclampsia marker genes *FLT1* and *FSTL3*. Several studies have also reported increased expression of other genes in this cluster in preeclampsia and intrauterine growth restriction, including *NDRG1*, *HK2*, and *TET3* (76–80), but none of these studies reported specifically on whether this increased expression was specific to one sex or shared in both sexes. Cluster 6 genes showed similar expression in male and female placentas from normotensive pregnancies, but changed in different directions in hypertensive pregnancies, showing lower expression in male placentas from hypertensive pregnancies and higher expression in female placentas from hypertensive pregnancies. This cluster did not show any significant enrichment for specific biological processes, but several of these genes showed upregulation in female placentas from uncomplicated first trimester pregnancies: *JPX*, *LINC00630*, *OFD1*, and *BRCC3* (81). Several have also been shown to be upregulated in the placenta in preeclamptic pregnancies: *NPFFR2* (82), *XIAP* (83), and *DLD* (84). The largest cluster of genes was cluster 4, suggesting that there are many genes that have lower expression in male placentas in normotensive pregnancies than any of the other placenta sample groups in our study. This cluster contained many known placental markers of preeclampsia, including *ENG*, *INHA*, *INHBA*, *PAPPA2*, *SLCO2A1*, *KRT19*, and *SPAG4*. Collectively, these gene expression patterns demonstrated that there are genes that show differences between placentas from pregnancies with different hypertension status and that those were overlaid onto sex specific differences in placental gene expression.

**Figure 5.**
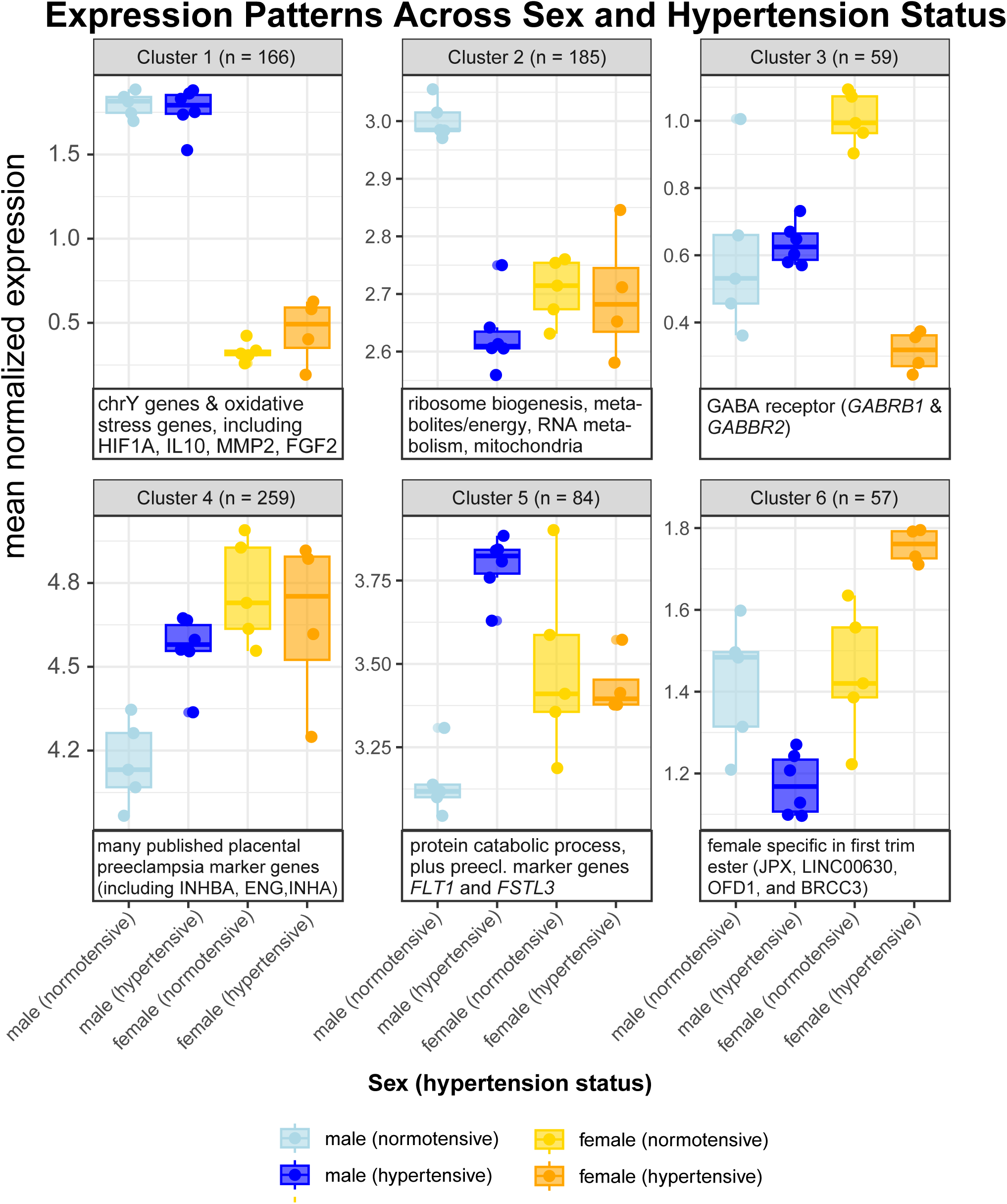
Differential expression patterns across placentas grouped by sex and pregnancy hypertension status. Mean normalized expression across top 6 clusters for genes that show nominal significance (p-value < 0.01) in at least one comparison between groups based on sex and pregnancy hypertension status.

### Sex-stratified gene expression signatures overlap with preeclampsia cohorts

To assess the reproducibility of our results, we identified overlap in sex-straified gene expression profiles across the three independent placenta studies that include samples from hypertensive disorders of pregnancy. We identified two published datasets that used RNA sequencing to measure gene expression in the placenta where fetal sex was annotated for each placenta sample. The first analyzed placental gene expression data from late-onset preeclampsia with typical features compared to placentas from uncomplicated term pregnancies in a pregnancy cohort from Adelaide, South Australia, Australia, where differences in placental gene expression were found in only male-bearing preeclamptic pregnancies (16). The second measured gene expression from placenta in a cohort with preeclampsia selected with severe features from San Diego, California, USA, which we reanalyzed as sex-stratified differential expression profiles of preeclampsia using matched analytical protocols (34). To maximize sensitivity to overlap in different degrees of differential expression across data with different numbers of samples per placenta, we used the threshold-free approach of rank-rank hypergeometric analysis to compare full expression profiles created by ranking all genes by log_2_-transformed fold change in gene expression in placenta from hypertensive versus normotensive pregnancies separately in male placentas and female placentas (85). The sex-stratified placental gene expression profile of hypertensive pregnancies were compared pairwise across the three data sets.

We observed the highest overlap between our dataset and that of Horii *et al*. that comprised pregnancies with preeclampsia with severe features. In the males, the overlapping signal was dominated by genes that had higher expression in placenta from hypertensive pregnancies in both datasets (Figure 6, Supplemental Figure 5). The overlapping gene set was enriched for genes involved in pregnancy (including several prostaglandin genes), inflammatory and immune responses, transport of biomolecules, and regulation of body fluid levels. In females, overlap was observed on both sides of the ranked lists (genes with both higher and lower expression in hypertensive/preeclamptic pregnancy placentas compared to normotensive pregnancies). Genes with higher expression in female placentas from hypertensive pregnancies were enriched for cell adhesion including *ITGA* integrin genes, extracellular matrix including *MMP* and *ADAMTS* metalloproteases, and muscle contraction related genes including *CACN* calcium channel genes. Genes decreased in females in both datasets were enriched for mitochondrial ATP synthesis and gland development related genes. Genes annotated as related to gland development include *EGFR*, which previous studies have shown its deficiency causes placental abnormalities due to reduced trophoblast proliferation when knocked out (86). This enrichment also includes *CDH1* (e-cadherin), which is downregulated to permit invasion of extravillous cytotrophoblasts into maternal uterine tissue and loss of cadherin was shown in cytotrophoblasts in preeclamptic pregnancies (87,88). In males, genes with higher expression in hypertensive (preeclamptic) pregnancies in both our and the Horii *et al.* datasets were enriched for genes involved in pregnancy, T cell proliferation, and negative regulation of viral genome replication. Pregnancy genes with increased expression in male placentas from hypertensive pregnancies include several pregnancy-specific glycoprotein genes (*PSG* genes), which are secreted from syncytiotrophoblasts increasingly more as pregnancy continues and have immunoregulatory, pro-angiogenic, and anti-platelet functions at the feto-maternal interface (89,90). Genetic variants in *PSG* genes have been reported to confer risk for preeclampsia (91), but our study suggests that these genes might be more involved in pregnancy complications including gestational hypertension in male-bearing pregnancies. Genes decreased in male placentas from hypertensive pregnancy across our and the Horii *et al.* data set included genes involved in metabolism of small molecules and nucleotides. This could indicate reduced cell proliferation or cell growth in the placenta of hypertensive pregnancies in males.

**Figure 6.**
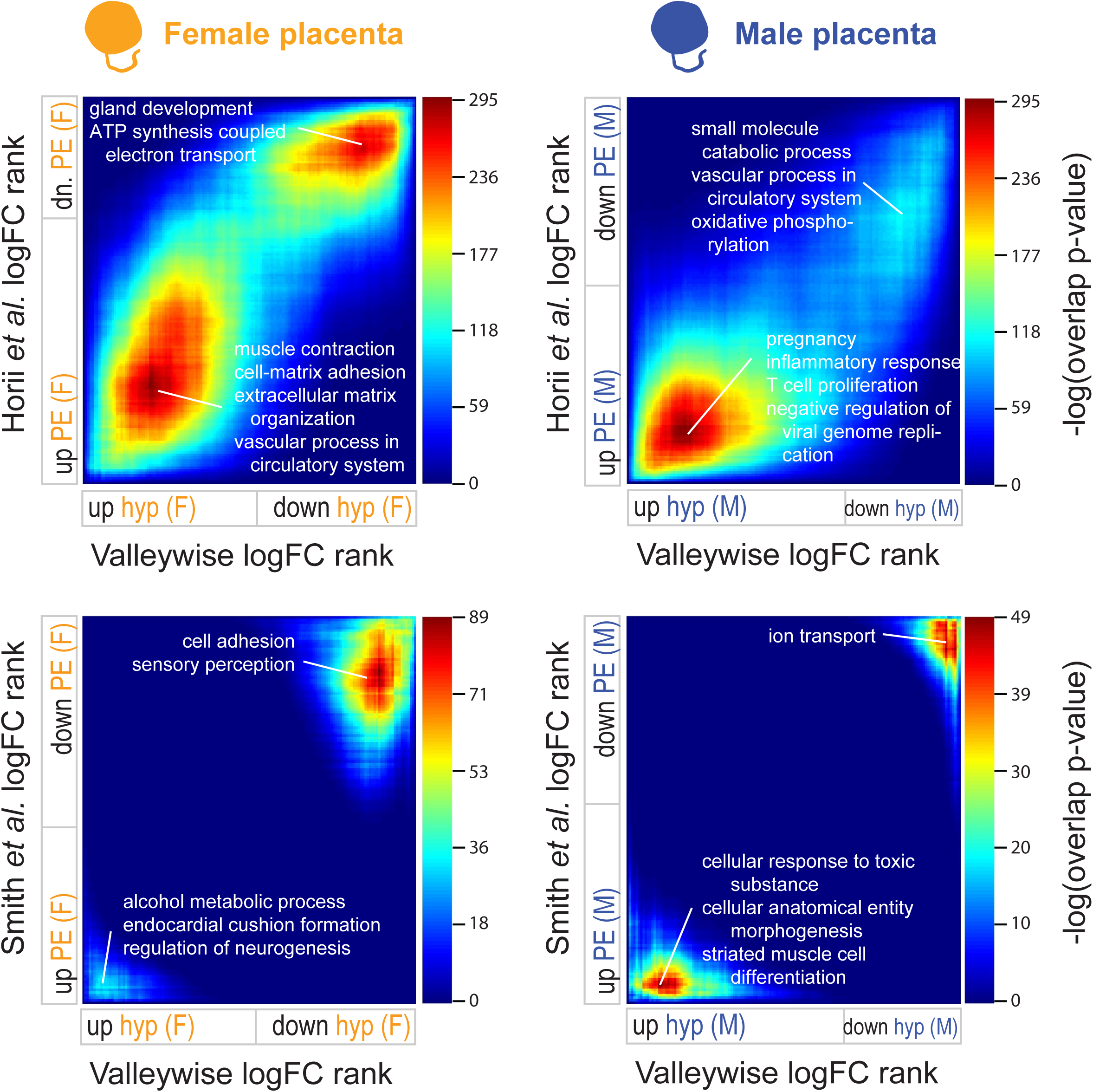
Overlap between sex-stratified differentially expressed genes in late-onset gestational hypertension with two published preeclampsia datasets. Rank-rank hypergeometric overlap (RRHO) maps created using genes ranked on moderated t-statistic between placentas from hypertensive/preeclamptic pregnancies compared to normotensive/control pregnancies are shown. Color bar indicates statistical significance of overlap; red indicates most significant overlap across the entire map. Significantly enriched biological processes in most significantly overlapping genes from the regions of the map that correspond to genes upregulated in both datasets and downregulated in both datasets.

When comparing both late-onset cohorts with typical features, ours and that of Smith *et al.*, overlap in male placentas was observed in genes that had higher and lower expression in placentas from hypertensive pregnancies, while in female placentas, overlap was mostly in genes that had lower expression in hypertensive pregnancies in both datasets (Figure 6, Supplemental Figure 6). In males, genes with higher expression in both late-onset studies were enriched for cellular response to toxic substances and cellular morphogenesis. Genes decreasing in males in both late-onset studies were enriched for anion transport including several GABA receptor genes. More significant overlap signal was observed in female placentas in genes that had lower expression in hypertensive versus normotensive pregnancies in both late-onset datasets; these overlapping genes were enriched for cell adhesion including several cadherin genes which have been shown to affect cytotrophoblast invasion in preeclampsia (92–94). The Smith *et al*. dataset also showed less significant overlap when compared to the Horii *et al*. dataset, so this may indicate that the Smith dataset includes more population specific gene expression differences between placentas from preeclamptic pregnancies versus those without preeclampsia(Supplemental Figure 4). Using these three datasets, we identified genes that were consistent across sex-stratified placental gene expression profiles of hypertensive disorders of pregnancy.

### Genes with high intraplacental variation involve processes important to the establishment of a proper feto-maternal interface

Having sampled each placenta in our study at 4 locations across the placental disc, we further assessed the intraplacental variation for each gene. As gene expression may vary more at different gene expression levels, we fit each gene’s variance in expression across the 4 samples in each placenta to the mean expression, then examined the residual variance such that genes with a large positive residual variance have much more variation than would be expected given their mean expression level. Median intraplacental variation across all placentas was normally distributed with a slight right skew due to more genes having high intraplacental variation compared to genes with conserved multiregional expression (Supplemental Figure 7A). Comparing median intraplacental variation in placentas grouped by fetal sex or hypertension status, intraplacental variation was highly correlated showing that intraplacental variation is consistent across sample groups (Supplemental Figure 7 B,C). Looking at the genes with the highest residual intraplacental variance across all the placentas in our dataset (Figure 7A), we show two patterns. The expression of some genes such as *IGFBP1* showed the full range of expression across the four sample sites. Some genes such as *DLG5* and *SQSTM1* show more of a bimodal distribution of expression with some sites with high expression and others with low to no expression. Genes that showed the least intraplacental variation such as *NEK9*, *TBRG1*, and *HNRNPU* showed similar expression across the four sites. Genes with low intraplacental variation were involved in essential cell functions needed for cell viability: *NEK9* is a kinase that regulates cell cycle progression (95), *TBRG1* is involved in maintaining chromosomal stability and regulating cell cycle arrest (96), and *HNRNPU* is an RNA-binding protein involved in transcription and translation (97). Genes with higher intraplacental variation may need more careful and extensive sampling to fully capture gene expression differences in placenta studies.

**Figure 7.**
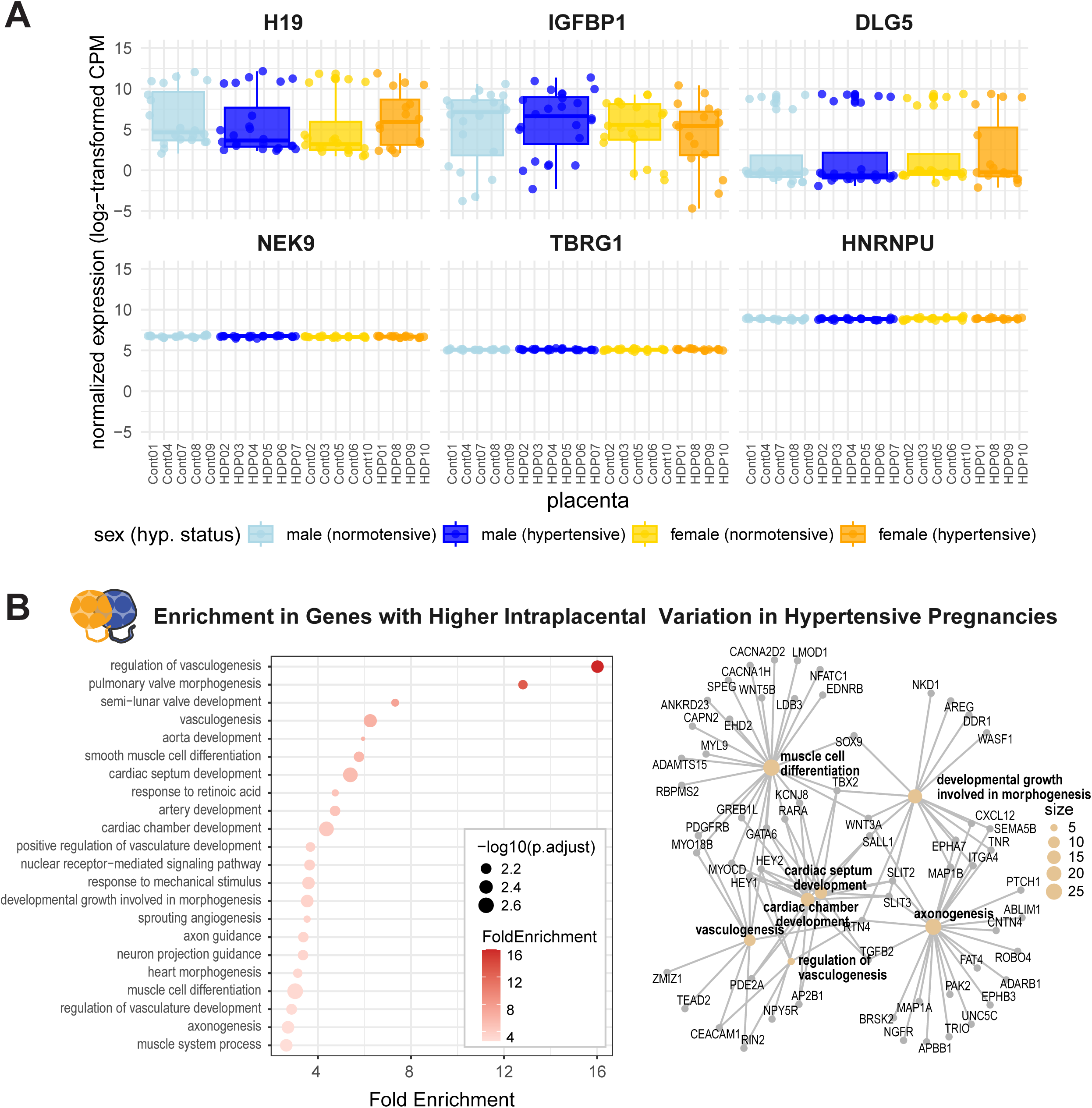
Genes with highest and lowest intraplacental variation across all placentas. (**A**) Boxplots showing the expression of the top 3 genes with highest and lowest intraplacental variation across all placentas. (**B**) Dot plot and gene connectivity map showing significantly enriched biological processes in genes that had higher intraplacental variation in placentas from hypertensive pregnancies (t-test p-value < 0.05).

Gene set enrichment analysis from ranking the genes by median intraplacental variation across all placentas in our study identified processes that were highly variable or highly conserved across the placental disc (Figure 8B). Genes with the highest intraplacental variation were most enriched for epithelial-to-mesenchymal transition (EMT). EMT is central to the differentiation of cytotrophoblasts into invasive extravillous trophoblasts (98). Highly variable genes were also enriched for coagulation and inflammatory response; tight regulation of coagulation in trophoblasts that affects genes involved in growth and signaling has been shown to be necessary for proper placenta function in mice and inflammatory signals can disrupt this regulation (99). Highly variable genes were enriched for estrogen response, which is involved in development of placental vasculature and remodelling of the spiral arteries in the maternal uterine decidua (100). Apoptosis was enriched; apoptosis has been observed at high levels in syncytiotrophoblasts at late gestation in tertiary placental villi as parts of the apoptosis machinery is used when cytotrophoblasts fuse into the syncytium (101). Apoptosis has been shown to be altered in several pregnancy conditions associated with abnormal placenta morphology; it is increased in conditions affecting late pregnancy including preeclampsia (102–104). Genes sets involving hypoxia, glycolysis, and reactive oxygen species were also enriched in genes with high variation. Studies have suggested that preeclampsia involves oxidative stress in trophoblast which induces apoptosis and trophoblast turnover which leads to excessive release of cytotoxic factors into maternal blood circulation. Disruption of blood delivery causes local hypoxia which can lead to increased production of reactive oxygen species and increased glycolysis (105). Increased intraplacental variation in genes involved in this process could explain or reflect different stages of this process at different locations in the placentas in our study. Genes with the lowest observed intraplacental variation were significantly enriched for gene sets involving targets of Myc transcription factor, DNA repair, and protein secretion. The Myc targets gene set reflects genes involved in cell proliferation (106); when combined with DNA repair and transport and processing of proteins, genes with low intraplacental variation were involved in cellular functions essential for viability and proper function of cells. All in all, genes with high intraplacental variability across all the placentas in our dataset were enriched for many functions involved in the establishment of the materno-fetal interface and variation in their expression may reflect variation in progenitor trophoblast stem cells in different lobes across the placental disc.

**Figure 8.**
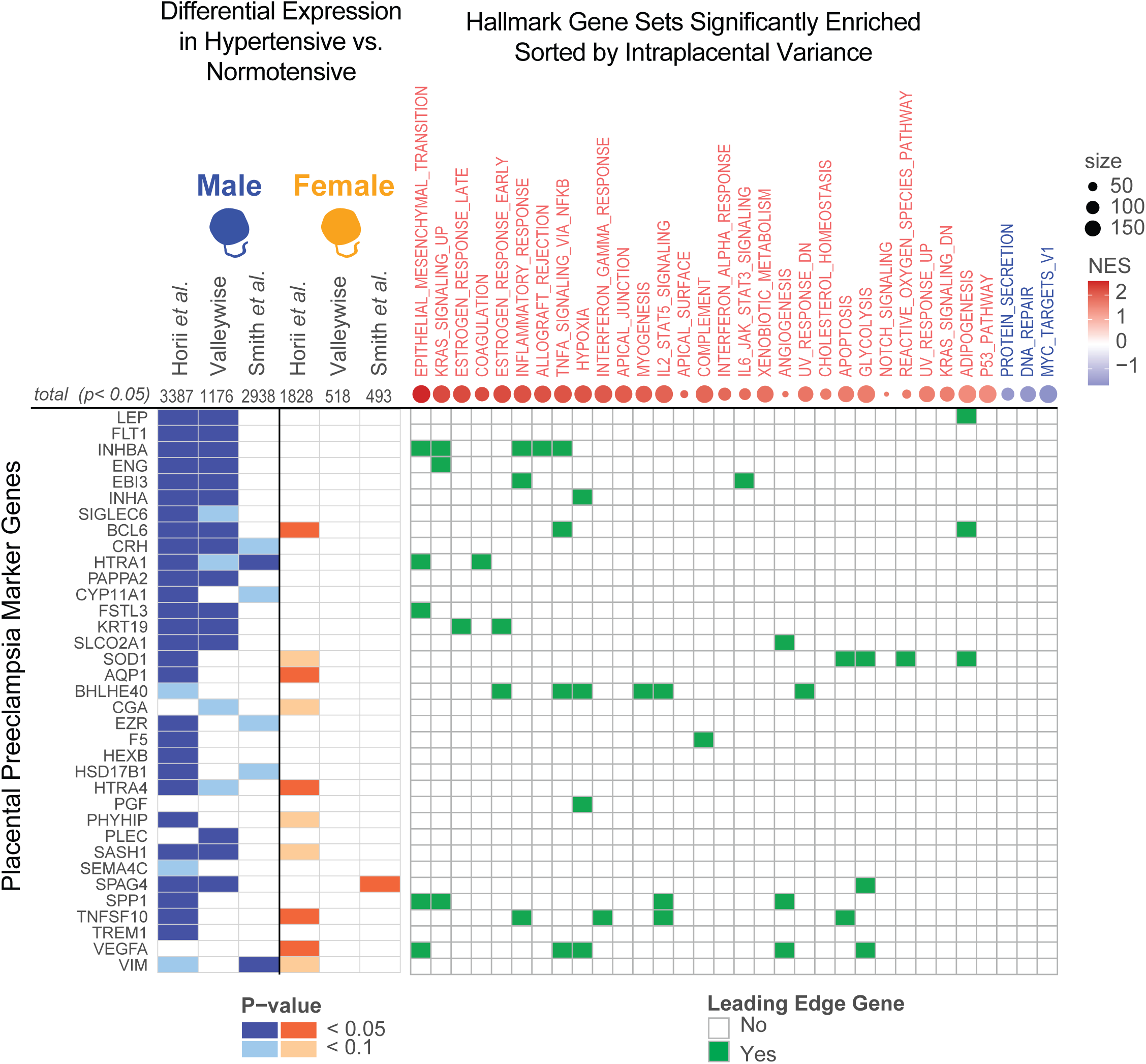
Published placental preeclampsia marker genes are more commonly differentially expressed in males and part of the most variably expressed biological process gene sets. Placental preeclampsia marker genes are sorted in order from most widely reported at the top to least widely reported at the bottom. Sex-stratified differential gene expression analysis was run with three datasets studying hypertensive disorders of pregnancy; total number of genes that had p-value < 0.05 are listed for each sex-stratified profile. Each marker gene is marked according to differential expression in each profile, with blue representing differential expression in male placentas and orange in female placentas, with darker shading indicating differential expression p-value < 0.05 and lighter shading indicating p-value < 0.1. Marker genes were also labeled for leading edge membership in gene sets that were significantly enriched when sorting genes from most to least median intraplacental variation across all placentas; normalized enrichment scores (NES) are shown for all Hallmark gene sets with adjusted enrichment p-value < 0.05 and size of the enriched gene set is indicated by point size.

To determine if this intraplacental variation in gene expression is affected by gestational hypertension, we examined genes that had significantly higher intraplacental variation in placentas from hypertensive pregnancies compared to normotensive pregnancies. As intraplacental variation within male and female placentas were well correlated (Supplemental Figure 7 B,C), we compared variation in all placentas from hypertensive pregnancies to all placentas from normotensive pregnancies. 521 genes had significantly higher intraplacental variation (t-test p-value < 0.05) in placenta from hypertensive pregnancies and these genes were significantly enriched for several aspects of blood vessel formation (sprouting angiogenesis, vasculogenesis, artery development), muscle cell differentiation, response to mechanical stimulus, and axon guidance (Figure 7B). Higher variation in gene expression for genes involved in these processes could indicate that (1) at some locations in the placenta, low expression of these genes may reflect a lower capacity of the placenta to have invaded into the uterine tissue, remodel uterine blood vessels, and established proper feto-maternal exchange in hypertensive pregnancies at some point during gestation or (2) at some locations, there will be high expression of these genes, which could indicate that a compensatory mechanism had taken place earlier in gestation so that the placenta tried to increase angiogenesis, morphogenesis, and metabolic activity in response to damaged systemic endothelial function in hypertensive pregnancies. Genes with increased variation in placenta from hypertensive pregnancies may reflect the mechanisms by which the placenta compensates in response to hypertension during pregnancy.

### Frequently reported preeclampsia marker genes were differentially expressed in males more than in females and were highly variable across the placental disc

As fetal sex is not always analyzed in the context of placental gene expression and pregnancy complications, we sought to determine whether genes previously reported to be differentially expressed in placenta from preeclamptic pregnancies were differentially expressed in both male-and female-bearing pregnancies or specifically in placenta from one fetal sex. Genes that were previously reported to have altered placental gene expression in preeclamptic pregnancies (107) showed more differential expression in male placentas than female placentas in our dataset as well as the Horii *et al.* and Smith *et al.* preeclampsia data sets (Figure 8, Supplemental Table 10). The Horii *et al.* severe feature preeclampsia study had the most placental preeclampsia marker genes differentially expressed. Nearly all preeclampsia markers were differentially expressed in male placenta (differential expression p-value < 0.05), while only 5 out of 35 were differentially expressed at that level in female placentas. In our dataset, about half of the placenta preeclampsia markers were differentially expressed in males but none were differentially expressed in females. In the Smith *et al.* dataset, 2 marker genes were differentially expressed (p-value < 0.05) and 4 marker genes were approaching significance (p-value < 0.1) in male placentas, but only one marker gene, *SPAG4,* was differentially expressed in female placentas. These findings underscore the importance of considering sex as a biological variable, as several markers thought to be broad placental markers of preeclampsia were only differentially expressed in male placentas in various hypertensive disorders of pregnancy. This demonstrates that potential therapeutic targets in the placenta may have sex differences in efficacy for ameliorating gestational hypertension and preeclampsia.

In addition to differential expression of these placental preeclampsia marker genes, we examined their intraplacental variation. When compared to the intraplacental variation for all genes, known placental gene markers of preeclampsia had among the highest levels of intraplacental variation (Figure 8, Supplemental Figure 8). The marker gene reported in the most studies as differentially expressed in placenta from preeclamptic pregnancies versus normotensive control pregnancies, *LEP*, was in the 95th percentile or higher by intraplacental variation in all placentas in our cohort. *LEP*, *CRH*, and *IGFBP1* each showed a median intraplacental variation above the 99th percentile. 17 marker genes were above the 90th percentile for intraplacental variance in at least half of the placenta samples: *LEP*, *IGFBP1*, *CRH*, *RDH13*, *HTRA4*, *FSTL3*, *INHBA*, *TNFSF10*, *INHA*, *BCL6*, *HTRA1*, *SLCO2A1*, *CGA*, *CYP11A1*, *VEGFA*, *EBI3*, and *PGF* (Supplemental Figure 8, Supplemental Table 9). Many of these placenta preeclampsia marker genes were in the leading edge genes in gene sets significantly enriched in genes ranked by mean intraplacental variation (Figure 8), suggesting that widely reported preeclampsia markers are part of processes that are highly variable across the placenta and thus their expression may be especially sensitive to the often random location the placenta is sampled for gene expression analysis.

## Discussion

Using sex-balanced, multiregional collection of samples from placenta tissues in full-term pregnancies with and without late-onset gestational hypertension, we show that genes differentially expressed between placenta from hypertensive and normotensive pregnancies differ by fetal sex. In our late-onset gestational hypertension dataset, as well as a published preeclampsia with severe features dataset, male placentas showed more significant differential expression in known preeclampsia marker genes, showing the importance of considering fetal sex when studying pregnancy complications. One of these genes, *FLT1*, has clinical significance as the ratio of soluble Flt-1 (sFlt-1) to PlGF in maternal blood typically evaluated between 23 and 35 weeks of gestation is used for preeclampsia prognosis and risk stratification (108). From early gestation in healthy pregnancies, the placenta produces placental growth factor (PlGF) which has angiogenic properties and low levels of anti-angiogenic sFlt-1. In preeclamptic pregnancies, expression of sFlt-1 increases significantly. A high sFlt-1/PlGF ratio is associated with higher risk for preeclampsia. We observe that *FLT1* was only differentially expressed in male placentas in our study of late-onset gestational hypertension (preeclampsia without severe effects) and in a previous study of preeclampsia with severe effects. Previous studies of sFlt-1/PlGF showed that in cases where a high ratio indicative of preeclampsia risk was observed, male-bearing pregnancies had higher ratios than female-bearing pregnancies (109). Also, sFLT1 overexpression in mouse models of preeclampsia leads to altered fetal liver development, specifically in male offspring (110). Early-onset preeclampsia has instead been associated with higher sFlt-1/PlGF ratio in female-bearing pregnancies than in male-bearing pregnancies (111,112). Taken together, gene expression differences in the placenta that have been broadly attributed to pregnancy complications may be unique to one sex and sex-stratified studies across many diverse populations could help to detail mechanisms underlying these sex differences which may inform differences in incidence and treatment of pregnancy complications.

We used a threshold-free approach to look for genes that were overlapping between sex-stratified placental gene expression profiles of preeclampsia or gestational hypertension from different studies. This allowed us to identify gene expression changes that were consistent across two or more datasets in genes that may not have been differentially expressed to a degree which would have allowed us to detect them with standard differential expression thresholds. For example, ceruloplasmin (*CP*) was a gene found to be significantly differentially expressed in male placentas between pregnancies with and without late-onset preeclampsia in the Smith *et al.* study (16) and was identified in the overlap with our study in male placentas. *CP* had 2.1 times higher expression on average in male placentas from hypertensive pregnancies compared to normotensive males in our dataset but this did not reach statistical significance, partly because *CP* expression was highly variable across the 4 samples in our placenta tissues. *CP* expression was in the 90th percentile by intraplacental variation in male placentas (and 95th in female placentas) so it is plausible that the Smith *et al.* study happened to take a single sample at locations in each placenta that had high expression of this gene in male placentas from preeclamptic pregnancies, but this might not have been captured if samples were taken from other locations in the placenta. Overall expression differences in placental gene expression can also be different across different population ancestry groups; additional studies with multiregional sampling and detailed descriptions of where placental tissue was sampled for gene expression analysis are needed to distinguish population-level expression differences from sampling-level differences. Genes with high intraplacental variation, specifically those with higher variation in placentas from hypertensive pregnancies, may reflect sites with different levels of angiogenesis, vascular remodeling, and placental invasion at the feto-maternal interface. Intraplacental variation might also reflect differences in the number of terminal villi at different sites allowing for differing levels of feto-placental exchange. Spatial transcriptomics would allow more detailed study of the cause of intraplacental variation in gene expression, including differences in the amount and character of placental cell types at different sites, cellular features such as density and shape of placental villi, exposures such as hypoxia, and tissue damage incurred during pregnancy or during delivery.

This study sheds light on many changes in gene expression in placenta at term with and without late-onset gestational hypertension, a time point which some studies have suggested does not reflect differences in the placenta but rather differences in maternal conditions such as obesity and metabolic disorders. All of the mothers in our study were Hispanic with admixed American ancestry and most did have elevated BMI, but we were able to detect significant gene expression differences in the placenta between different fetal sexes and hypertension status. In all three pregnancy cohorts that we analyzed which were from different parts of the world, there were more differentially expressed genes in male placentas with and without hypertensive disorders of pregnancy. We did see similarities and differences in the specific genes that were differentially expressed by these two features across three pregnancy cohorts. The mothers enrolled in this study were from a traditionally underrepresented community (Hispanic mothers). Importantly, this research provides a novel placenta genomics dataset that aimed to assess the impact of fetal sex in pregnancy complications. Additional studies of placenta from diverse populations with this, and other, pregnancy complications will enable the field to identify which genes are connected to which specific aspects of placental function in different sexes.

## Conclusions

In conclusion, sex differences in gene expression were observed in placenta from pregnancies with late-onset gestational hypertension compared to normotensive pregnancies (Figure 9). Late-onset gestational hypertension is a key symptom of late-onset preeclampsia, which has been understudied when compared to early-onset preeclampsia even though it is far more common. When comparing placenta from hypertensive pregnancies to normotensive pregnancies, placenta from each fetal sex showed a different profile of gene expression changes. Female placentas from hypertensive pregnancies showed lower expression of ribosome genes and higher expression of genes related to cell junction disassembly, response to angiotensin, PDGFRβ and TGFβ signaling, and T-cell and Toll-like receptor signaling. Male placentas showed lower expression of genes related to the mitochondria, amino acid metabolism, and DNA replication in addition to the ribosome, and higher expression of genes involved in the complement activation, humoral immune response, inflammatory response, and cell adhesion. Genes previously reported to be differentially expressed in the placenta in preeclamptic pregnancies were more differentially expressed in males, showing that previous findings on placental changes in preeclampsia might be sex-specific. Intraplacental variation in gene expression was consistent between male and female placentas. Genes with highly variable expression across the placental disc included known preeclampsia genes, which may contribute to inconsistencies between placenta differential expression studies of hypertensive disorders of pregnancy. Genes with higher variation in gene expression in placentas from hypertensive pregnancies across both sexes were enriched for genes associated with vasculogenesis, smooth muscle differentiation, response to mechanical stimuli, and axonogenesis, all of which are involved in the establishment of the fetomaternal interface, suggesting the variability in expression is a part of a placental response to or the cause of late-onset gestational hypertension.

**Figure 9.**
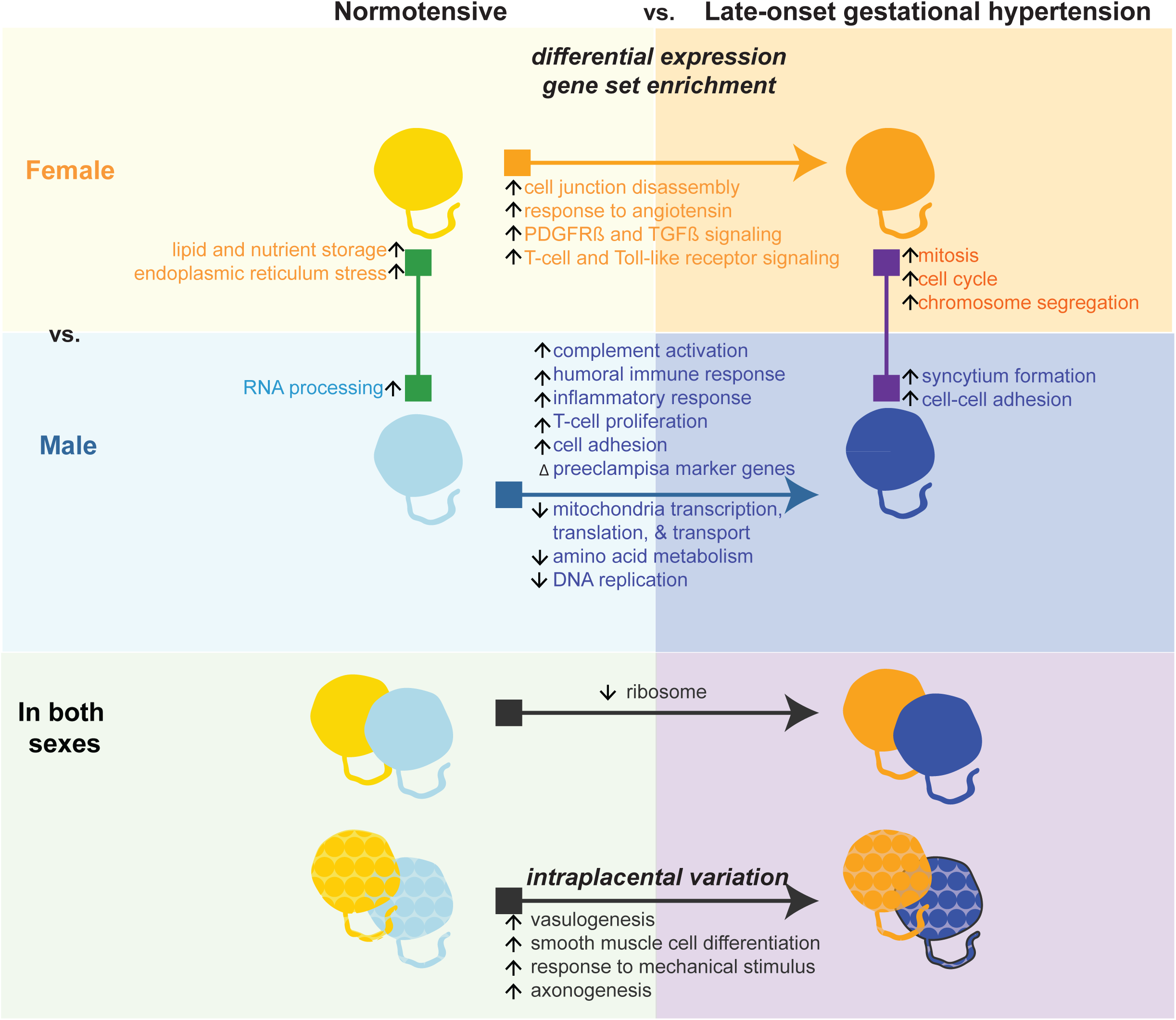
Summary of gene expression changes between placentas grouped by sex and pregnancy hypertension status. Summary of biological processes enriched in differential expression profiles between placentas from hypertensive versus normotensive pregnancies for each sex.

## Ethics approval and consent to participate

Ethics approval for use of human research subjects in this study and prospective collection of biological specimens for research purposes by noninvasive means was obtained from the Valleywise Health Medical Center Research Ethics Board (Protocol #2021-008, Approval Date 02/01/2021). Informed written consent was obtained from all study participants.

## Consent for publication

Not applicable

## Availability of data and materials

All raw genomics data collected for the placenta samples analyzed in this paper are available as controlled access through dbGap: phs003693.v1.p1. Code for analysis is available through GitHub: https://github.com/SexChrLab/Placenta_Multiomic_Multiregional/

## Authors’ Contributions

SP contributed to study design and data preparation, performed data analysis and interpretation, and drafted the manuscript. MW was lead investigator on the study, conceived of and oversaw all work and interpretation, procured research funding, edited the manuscript, and fostered collaborations needed for the work. AA expertly collected and stored placenta samples for sequencing. LN and DC provided clinical perspectives and contributed to the study design. MS and CR provided perspectives on placenta sample collection methods and genes involved in placenta biology and pregnancy complications. All authors read and provided critical feedback on the manuscript and approved the final version.

## Acknowledgements

We thank Anya Arthurs from Flinders University for final stage editing of the manuscript.

## Supplemental Figure Legends

**Supplemental Figure 1.**
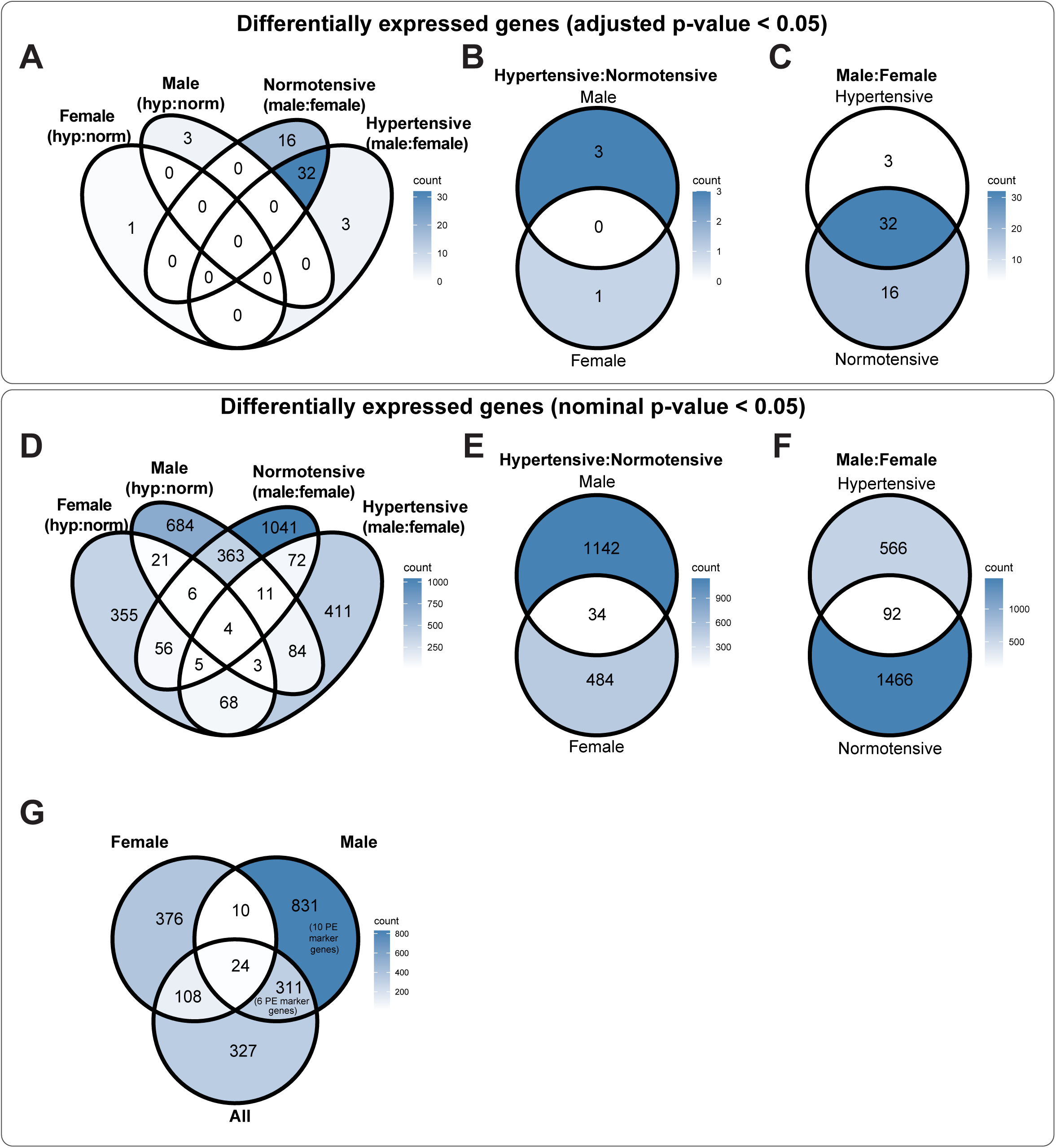
Overlap between differently expressed genes by sex and pregnancy hypertension status. Venn diagrams showing overlap between significant differentially expressed genes. Overlap in genes that were significant with (**A -C**) adjusted p-value < 0.05 and (**D-G**) nominal p-value < 0.05 are shown. (**G**) Venn diagram showing the number of differentially expressed genes in sex-stratified analysis of placentas from hypertensive pregnancies compared to non-sex stratified (all placentas from hypertensive versus all placentas from normotensive).

**Supplemental Figure 2.**
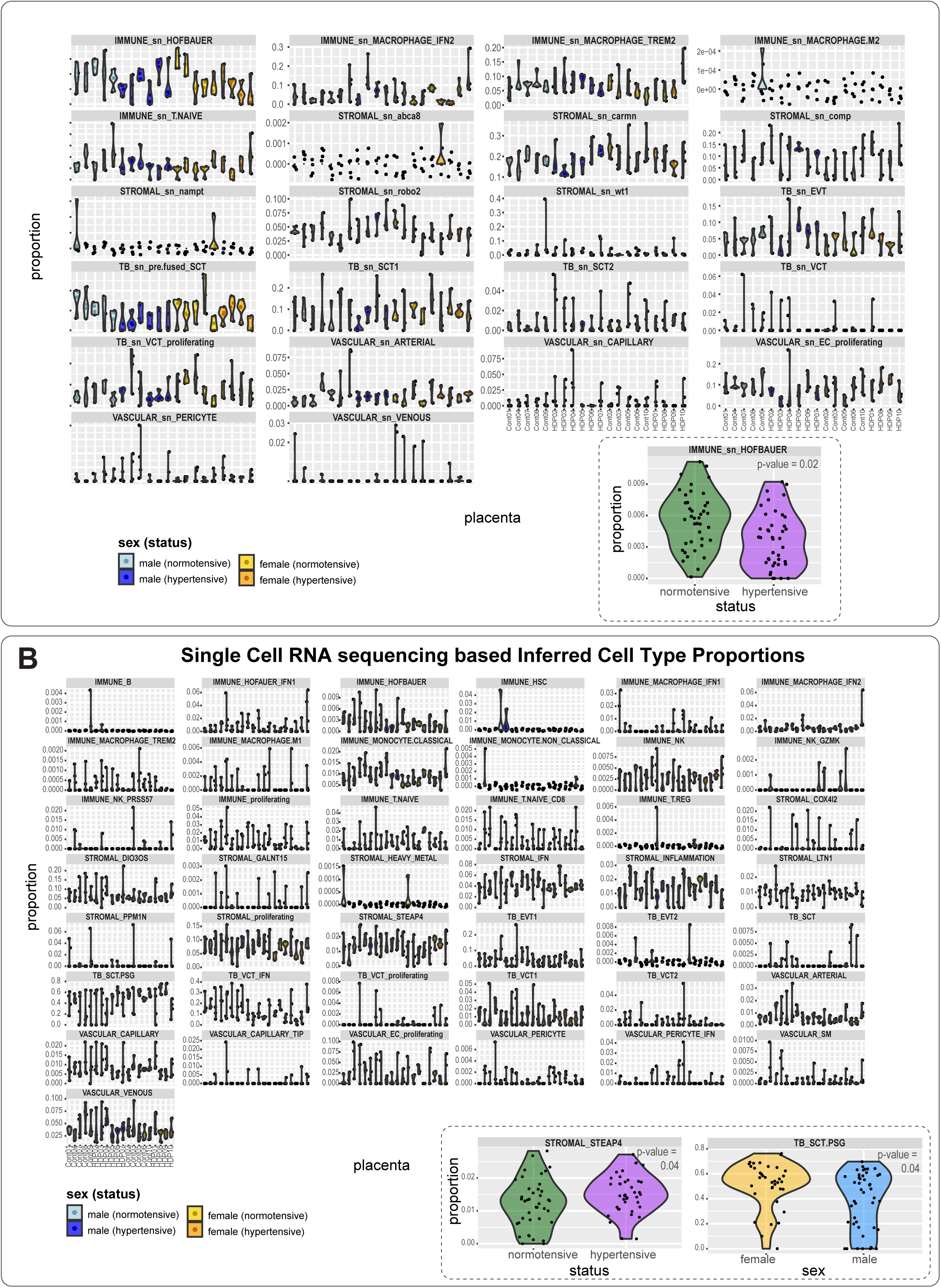
Inferred cell type proportions show no major differences between sex and pregnancy hypertension groups. Violin plots showing cell type proportions inferred using (**A**) single nucleus RNA sequencing and (**B**) single cell RNA sequencing. For each, inset violin plots show comparisons that have statistical significance when placentas with the same sex or same pregnancy hypertension status are combined.

**Supplemental Figure 3.**
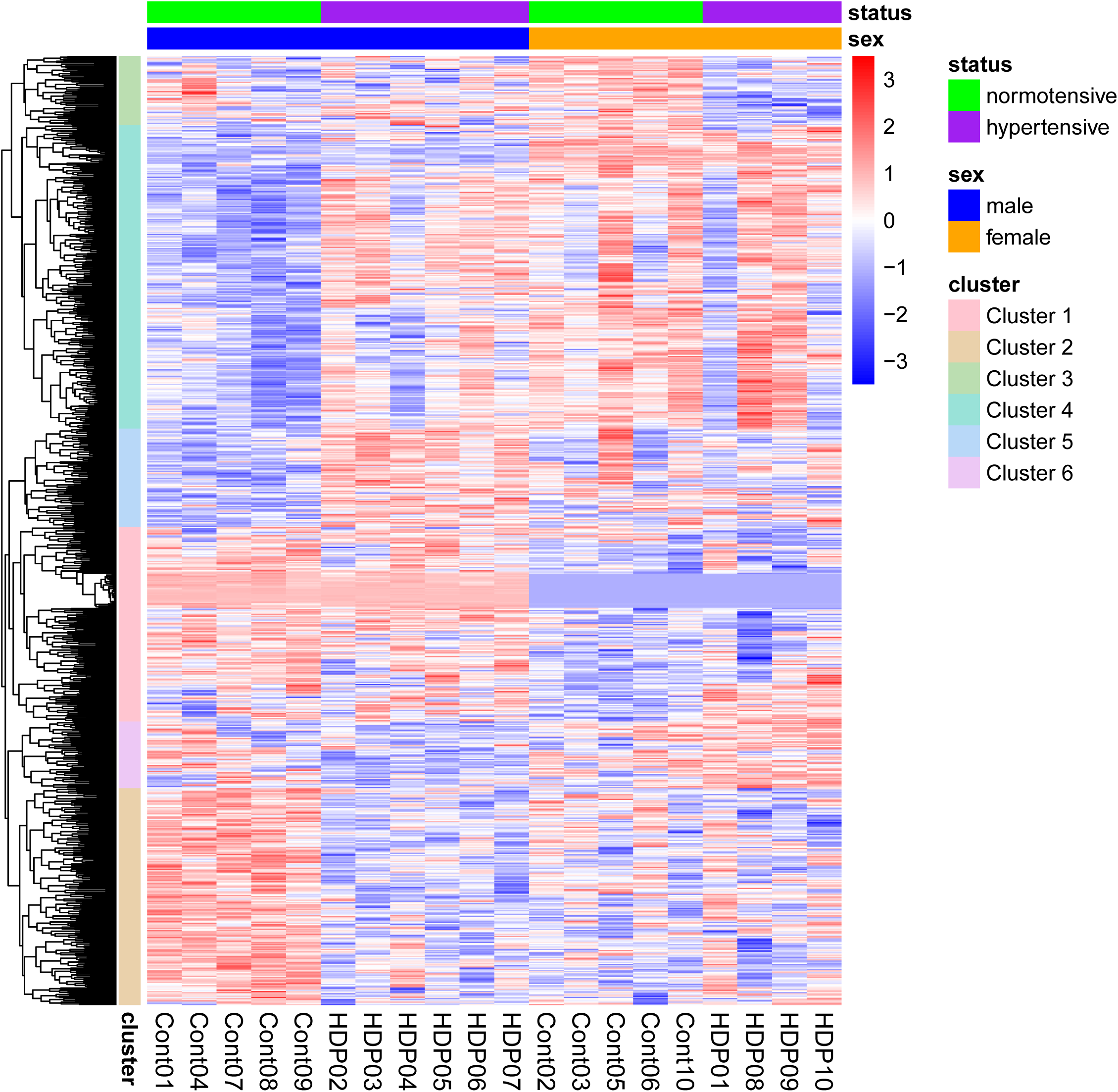
Gene expression patterns across placentas grouped by seed and pregnancy hypertension status. Heatmap showing clusters of genes with similar expression patterns in genes that had a nominal p-value value < 0.01 in at least one pairwise comparison.

**Supplemental Figure 4.**
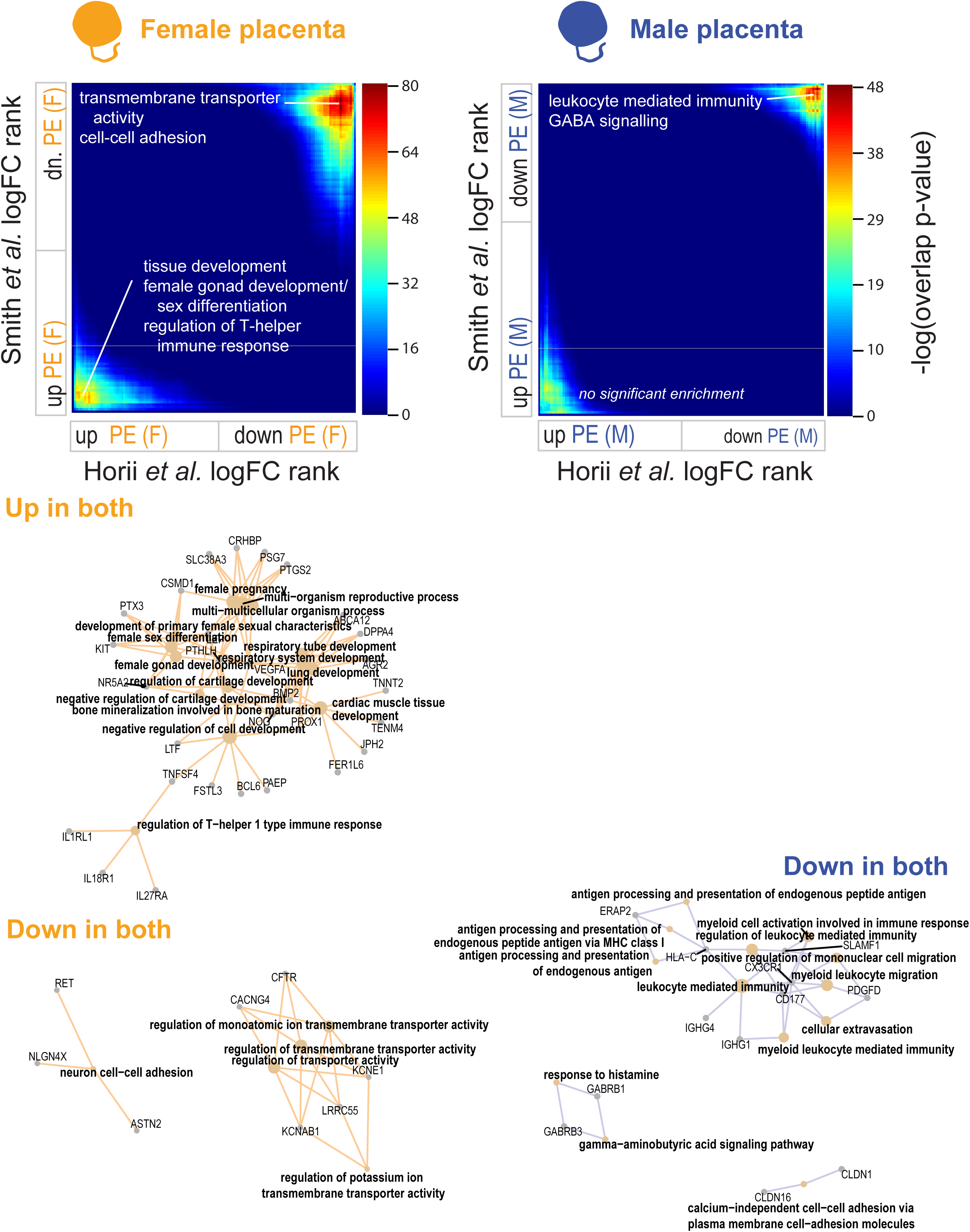
Overlap between sex-stratified differentially expressed genes in previously published preeclampsia datasets. Rank-rank hypergeometric overlap (RRHO) maps created using genes ranked on log_2_-transformed fold change between placentas from preeclamptic pregnancies compared to control pregnancies from Horii *et al.* and Smith *et al*. are shown. Color bar indicates statistical significance of overlap; red indicates most significant overlap across the entire map. Significantly enriched biological processes in most significantly overlapping genes from the regions of the map that correspond to genes upregulated in both datasets and downregulated in both datasets.

**Supplemental Figure 5.**
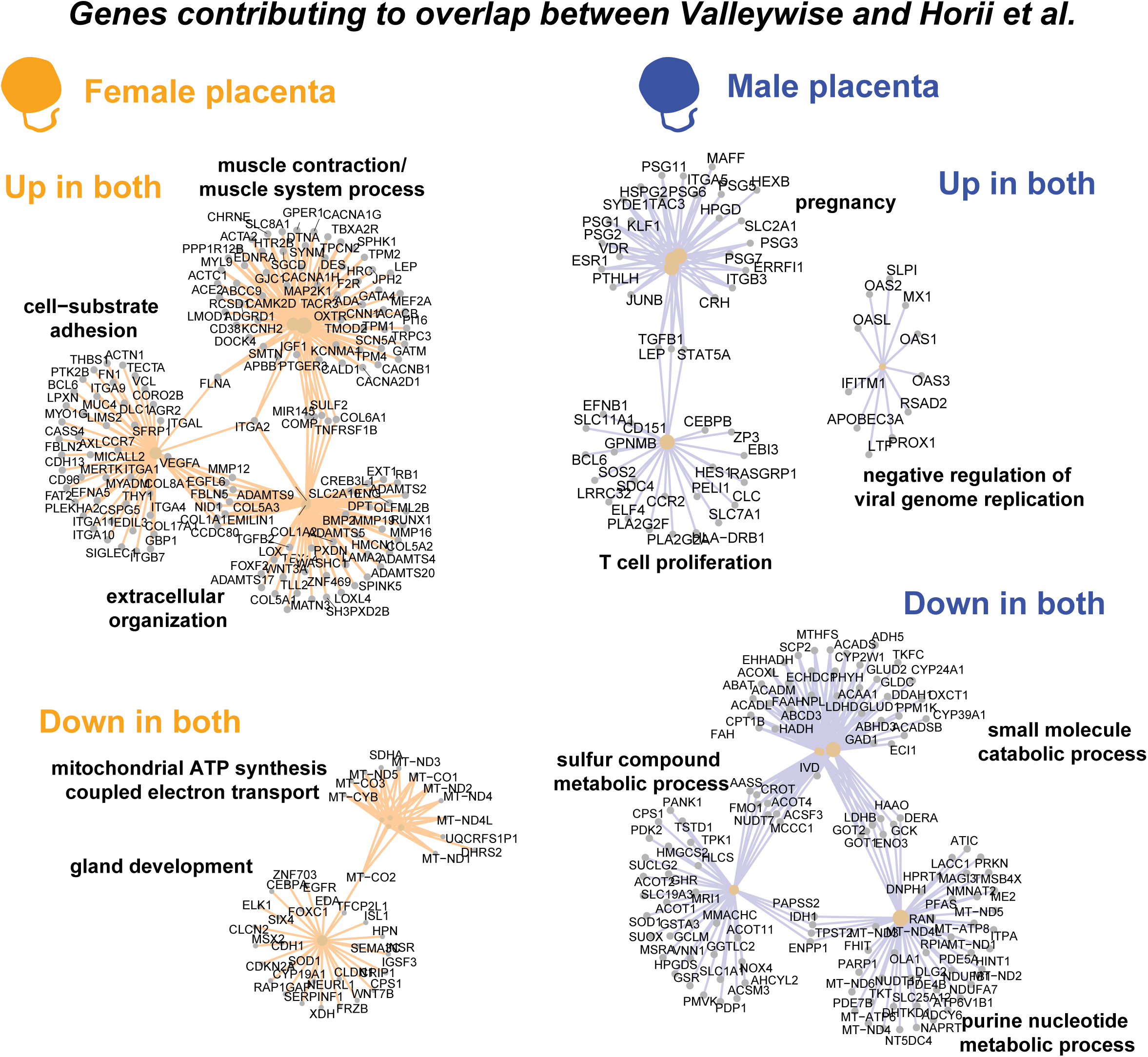
Genes contributing to overlap between Valleywise and Horii *et al*. Gene connectivity maps are shown for the top most significantly enriched biological processes for genes that have increased expression in both datasets and genes that have decreased expression in both datasets.

**Supplemental Figure 6.**
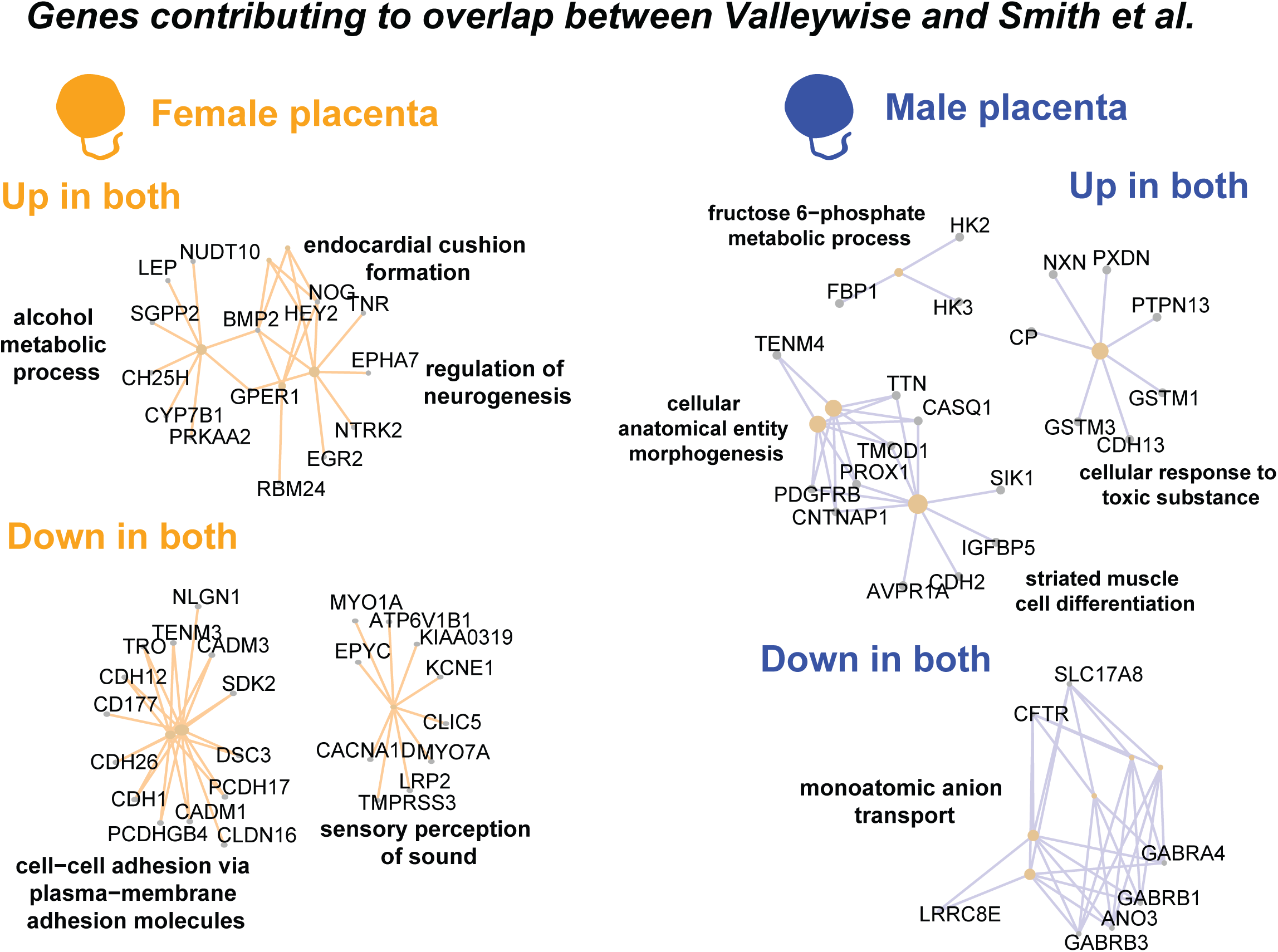
Genes contributing to overlap between Valleywise and Smith *et al*. Gene connectivity maps are shown for the top most significantly enriched biological processes for genes that have increased expression in both datasets and genes that have decreased expression in both datasets.

**Supplemental Figure 7.**
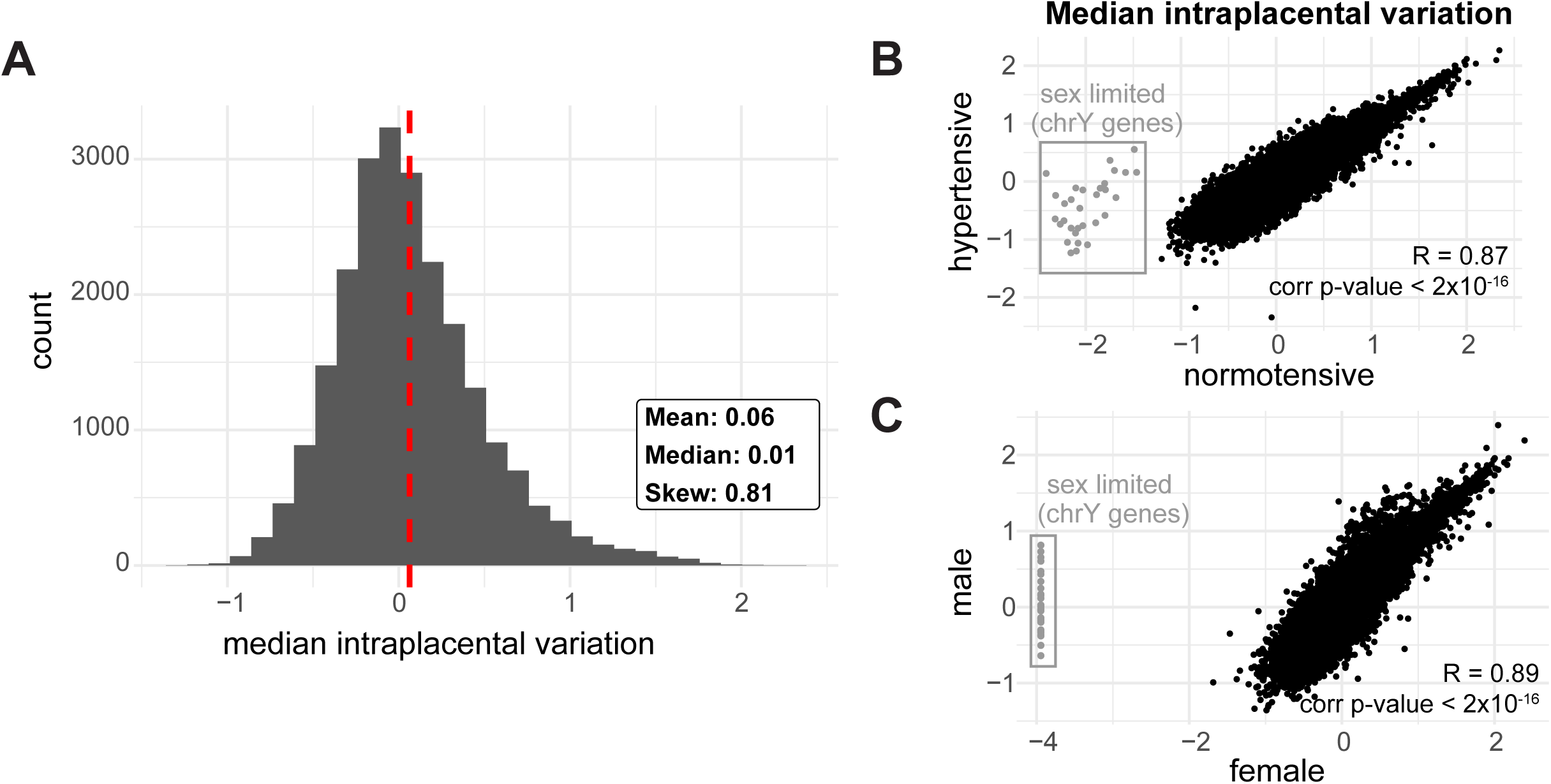
Intraplacental variance is consistent across sex and pregnancy hypertension status. (**A**) Histogram showing the range in median intraplacental variance across all placentas in the study; red dashed line labels the mean. (**B,C**) Scatter plots showing the correlation between median intraplacental variation across after splitting by (**B**) sex or (**C**) pregnancy hypertension status.

**Supplemental Figure 8.**
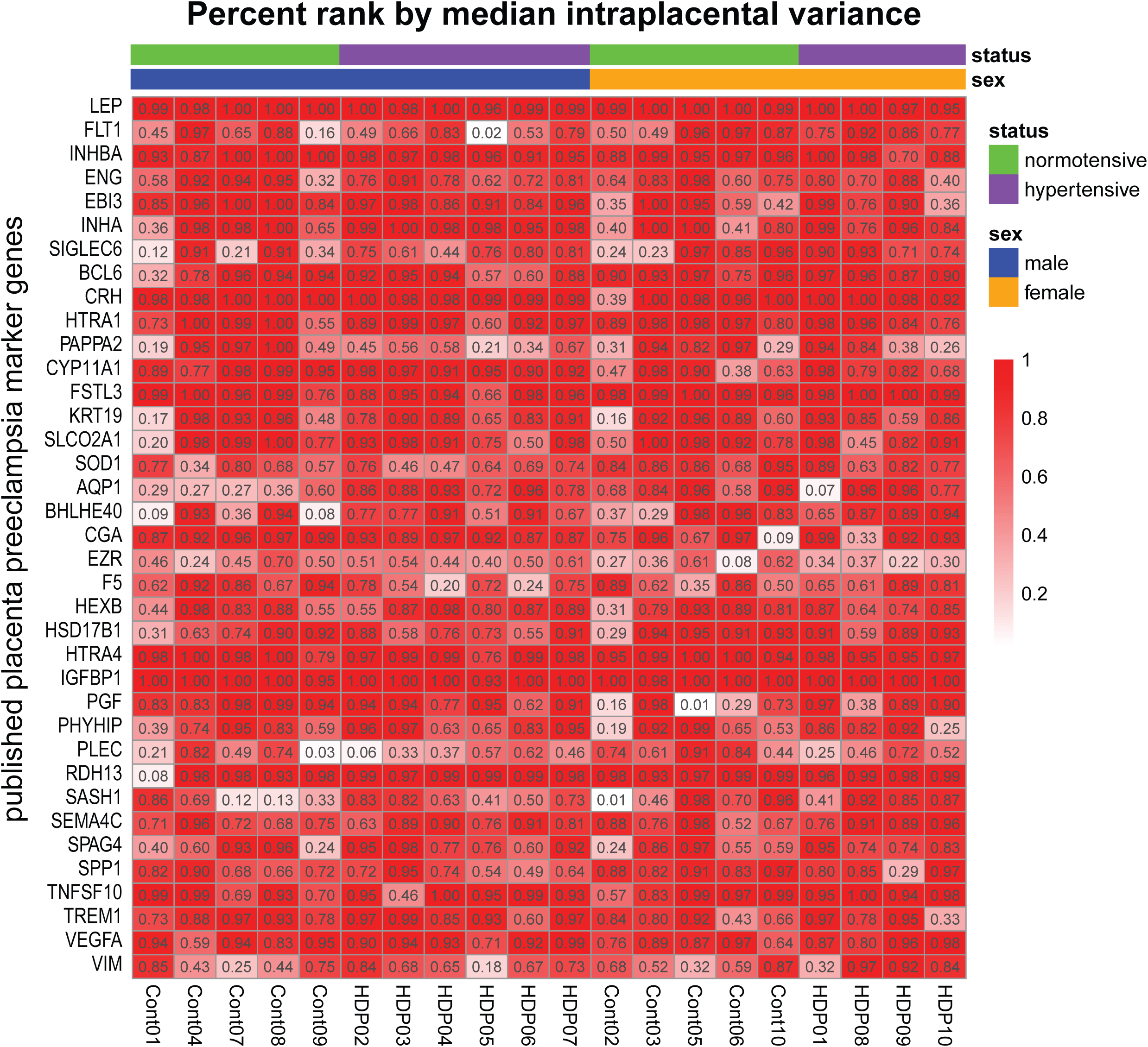
Published placenta preeclampsia genes are among the most variable in expression. Heatmap showing the percent rank for published preeclampsia marker genes when ordered by intraplacental variation. For each placenta, the mean of the intraplacental variance was taken across all 4 samples before percent rank was calculated.

## Table Legends

**Supplemental Table 1. Placenta and pregnancy sample metadata.** Information about fetal sex, delivery, characteristics of the parents, and clinically relevant information about the pregnancy and comorbidities.

**Supplemental Table 2. Pregnancy features by fetal sex and pregnancy hypertension status.** Appropriate statistical tests to show the difference in gestational age of the neonate at delivery, age and BMI of parents, gravidity, and parity grouping by fetal sex and pregnancy hypertension status.

**Supplemental Table 3. All placentas show Admixed American ancestry.** Variant calling summary statistics, genetic ancestry prediction, and principle components using exome sequencing for each placenta.

**Supplemental Table 4. No indication of chromosomal abnormalities in placenta samples**. Chromosome copy number estimates using exome sequencing for all autosomes, chromosome X, and chromosome Y.

**Supplemental Table 5. Differential expression results for male versus female placentas stratifying by pregnancy hypertension status.**

Tab 1– Differential gene expression results for all genes in normotensive pregnancies.

Tab 2– Differential gene expression results for all genes in hypertensive pregnancies.

Tab 3– Gene set enrichment analysis results in normotensive pregnancies.

Tab 4– Gene set enrichment analysis results in hypertensive pregnancies.

**Supplemental Table 6. Differential expression results for placentas from hypertensive versus normotensive pregnancies stratifying by fetal sex.**

Tab 1– Differential gene expression results for all genes in males.

Tab 2– Differential gene expression results for all genes in females.

Tab 3– Gene set enrichment analysis results in males

Tab 4– Gene set enrichment analysis results in females.

**Supplemental Table 7. Cluster assignments for significantly differentially expressed genes.**

Tab 1– Cluster assignment from hierarchical clustering of genes that have a nominal p-value < 0.01 in at least one pair of comparisons based on sex or pregnancy hypertension status.

Tab 2– Significantly enriched biological processes for each cluster.

**Supplemental Table 8. Intraplacental variation for all genes in all placentas.**

Tab 1– Intraplacental variance for all genes in all placentas with t-test p-value measuring the differential variance between placentas from hypertensive versus normotensive pregnancies and the median variance across all placentas.

Tab 2– Gene set enrichment analysis results ranking genes by decreasing intraplacental variance.

Tab 3– Significantly enriched biological processes in genes that had significantly higher intraplacental variance in placentas from hypertensive pregnancies.

**Supplemental Table 9. Differential expression of placenta preeclampsia marker genes in our dataset and two published preeclampsia data sets.** Log2-transformed fold change and differential expression p-values comparing placentas from hypertensive/preeclamptic pregnancies versus normotensive/control pregnancies for each sex in Valleywise, Horii *et al*., and Smith *et al*. datasets.

**Supplemental Table 10. Inferred cell type proportions.**

Tab 1– Single cell RNA sequencing based cell type deconvolution

Tab 2– Statistical significance of differences in cell type proportion inferred by single cell RNA sequencing based reference profiles grouped by sex and pregnancy hypertension status (4 groups)

Tab 3– Statistical significance of differences in cell type proportion inferred by single cell RNA sequencing based reference profiles grouped by pregnancy hypertension status (2 groups)

Tab 4– Statistical significance of differences in cell type proportion inferred by single cell RNA sequencing based reference profiles grouped by sex (2 groups)

Tab 5– Single nucleus RNA sequencing based cell type deconvolution

Tab 6– Statistical significance of differences in cell type proportion inferred by single nucleus RNA sequencing based reference profiles grouped by sex and pregnancy hypertension status (4 groups)

Tab 7– Statistical significance of differences in cell type proportion inferred by single nucleus RNA sequencing based reference profiles grouped by pregnancy hypertension status (2 groups)

Tab 8– Statistical significance of differences in cell type proportion inferred by single nucleus RNA sequencing based reference profiles grouped by sex (2 groups)

## References

1. Olney KC, Plaisier SB, Phung TN, Silasi M, Perley L, O’Bryan J, et al. Sex differences in early and term placenta are conserved in adult tissues. Biol Sex Differ. 2022 Dec 22;13(1):74. doi:10.1186/s13293-022-00470-y PubMed PMID: 36550527; PubMed Central PMCID: PMC9773522.

2. Oliva M, Muñoz-Aguirre M, Kim-Hellmuth S, Wucher V, Gewirtz ADH, Cotter DJ, et al. The impact of sex on gene expression across human tissues. Science. 2020 Sep 11;369(6509):eaba3066. doi:10.1126/science.aba3066

3. Derisoud E, Jiang H, Zhao A, Chavatte-Palmer P, Deng Q. Revealing the molecular landscape of human placenta: a systematic review and meta-analysis of single-cell RNA sequencing studies. Hum Reprod Update. 2024 Jul 1;30(4):410–41. doi:10.1093/humupd/dmae006

4. Braun AE, Mitchel OR, Gonzalez TL, Sun T, Flowers AE, Pisarska MD, et al. Sex at the interface: the origin and impact of sex differences in the developing human placenta. Biol Sex Differ. 2022 Sep 16;13(1):50. doi:10.1186/s13293-022-00459-7 PubMed PMID: 36114567; PubMed Central PMCID: PMC9482177.

5. Gilbert JS, Nijland MJ. Sex differences in the developmental origins of hypertension and cardiorenal disease. Am J Physiol-Regul Integr Comp Physiol. 2008 Dec;295(6):R1941–52. doi:10.1152/ajpregu.90724.2008

6. Gestational Hypertension and Preeclampsia: ACOG Practice Bulletin, Number 222. Obstet Gynecol. 2020 Jun;135(6):e237–60. doi:10.1097/AOG.0000000000003891

7. Chappell LC, Cluver CA, Kingdom J, Tong S. Pre-eclampsia. The Lancet. 2021 Jul;398(10297):341–54. doi:10.1016/S0140-6736(20)32335-7

8. Kariori M, Katsi V, Tsioufis C. Late vs. Early Preeclampsia. Int J Mol Sci. 2025 Nov 16;26(22):11091. doi:10.3390/ijms262211091 PubMed PMID: 41303573; PubMed Central PMCID: PMC12652060.

9. Lyall F, Robson SC, Bulmer JN. Spiral Artery Remodeling and Trophoblast Invasion in Preeclampsia and Fetal Growth Restriction: Relationship to Clinical Outcome. Hypertension. 2013 Dec;62(6):1046–54. doi:10.1161/HYPERTENSIONAHA.113.01892

10. Soto E, Romero R, Kusanovic JP, Ogge G, Hussein Y, Yeo L, et al. Late-onset preeclampsia is associated with an imbalance of angiogenic and anti-angiogenic factors in patients with and without placental lesions consistent with maternal underperfusion. J Matern-Fetal Neonatal Med Off J Eur Assoc Perinat Med Fed Asia Ocean Perinat Soc Int Soc Perinat Obstet. 2012 May;25(5):498–507. doi:10.3109/14767058.2011.591461 PubMed PMID: 21867402; PubMed Central PMCID: PMC3401571.

11. Lisonkova S, Joseph KS. Incidence of preeclampsia: risk factors and outcomes associated with early-versus late-onset disease. Am J Obstet Gynecol. 2013 Dec;209(6):544.e1–544.e12. doi:10.1016/j.ajog.2013.08.019

12. Baylis A, Zhou W, Menkhorst E, Dimitriadis E. Prediction and prevention of late-onset pre-eclampsia: a systematic review. Front Med. 2024;11:1459289. doi:10.3389/fmed.2024.1459289 PubMed PMID: 39640984; PubMed Central PMCID: PMC11617856.

13. Vatten LJ, Skjaerven R. Offspring sex and pregnancy outcome by length of gestation. Early Hum Dev. 2004 Jan;76(1):47–54. doi:10.1016/j.earlhumdev.2003.10.006 PubMed PMID: 14729162.

14. Tranquilli AL, Brown MA, Zeeman GG, Dekker G, Sibai BM. The definition of severe and early-onset preeclampsia. Statements from the International Society for the Study of Hypertension in Pregnancy (ISSHP). Pregnancy Hypertens. 2013 Jan;3(1):44–7. doi:10.1016/j.preghy.2012.11.001 PubMed PMID: 26105740.

15. Broere-Brown ZA, Adank MC, Benschop L, Tielemans M, Muka T, Gonçalves R, et al. Fetal sex and maternal pregnancy outcomes: a systematic review and meta-analysis. Biol Sex Differ. 2020 May 11;11(1):26. doi:10.1186/s13293-020-00299-3 PubMed PMID: 32393396; PubMed Central PMCID: PMC7216628.

16. Smith MD, Plaisier S, Breen J, Bogias KJ, Jankovic-Karasoulos T, McCullough D, et al. Sex-specific placental transcriptome alterations in late-onset preeclampsia reveal male-biased immune and metabolic dysregulation. Biol Sex Differ. 2025 Dec 24;17(1):8. doi:10.1186/s13293-025-00781-w PubMed PMID: 41444673; PubMed Central PMCID: PMC12809948.

17. Burton GJ, Sebire NJ, Myatt L, Tannetta D, Wang YL, Sadovsky Y, et al. Optimising sample collection for placental research. Placenta. 2014 Jan;35(1):9–22. doi:10.1016/j.placenta.2013.11.005

18. Avila L, Yuen RK, Diego-Alvarez D, Peñaherrera MS, Jiang R, Robinson WP. Evaluating DNA methylation and gene expression variability in the human term placenta. Placenta. 2010 Dec;31(12):1070–7. doi:10.1016/j.placenta.2010.09.011 PubMed PMID: 20947161.

19. Hughes DA, Kircher M, He Z, Guo S, Fairbrother GL, Moreno CS, et al. Evaluating intra- and inter-individual variation in the human placental transcriptome. Genome Biol. 2015 Mar 19;16(1):54. doi:10.1186/s13059-015-0627-z PubMed PMID: 25887593; PubMed Central PMCID: PMC4404591.

20. Mayhew TM. Changes in fetal capillaries during preplacental hypoxia: growth, shape remodelling and villous capillarization in placentae from high-altitude pregnancies. Placenta. 2003;24(2–3):191–8. doi:10.1053/plac.2002.0895 PubMed PMID: 12566246.

21. Chen CP, Aplin JD. Placental extracellular matrix: gene expression, deposition by placental fibroblasts and the effect of oxygen. Placenta. 2003 Apr;24(4):316–25. doi:10.1053/plac.2002.0904 PubMed PMID: 12657504.

22. FastQC: a quality control tool for high throughput sequence data [Internet]. 2010. Available from: http://www.bioinformatics.babraham.ac.uk/projects/fastqc/

23. Ewels P, Magnusson M, Lundin S, Käller M. MultiQC: summarize analysis results for multiple tools and samples in a single report. Bioinformatics. 2016 Oct 1;32(19):3047–8. doi:10.1093/bioinformatics/btw354 PubMed PMID: 27312411; PubMed Central PMCID: PMC5039924.

24. Bushnell B. BBMap: A Fast, Accurate, Splice-Aware Aligner. Annual Genomics of Energy & Environment Meeting. Walnut Creek, CA. (Lawrence Berkeley National Laboratory).

25. Olney KC, Brotman SM, Andrews JP, Valverde-Vesling VA, Wilson MA. Reference genome and transcriptome informed by the sex chromosome complement of the sample increase ability to detect sex differences in gene expression from RNA-Seq data. Biol Sex Differ. 2020 Jul 21;11(1):42. doi:10.1186/s13293-020-00312-9 PubMed PMID: 32693839; PubMed Central PMCID: PMC7374973.

26. Webster TH, Couse M, Grande BM, Karlins E, Phung TN, Richmond PA, et al. Identifying, understanding, and correcting technical artifacts on the sex chromosomes in next-generation sequencing data. GigaScience. 2019 Jul 1;8(7):giz074. doi:10.1093/gigascience/giz074

27. Kim D, Langmead B, Salzberg SL. HISAT: a fast spliced aligner with low memory requirements. Nat Methods. 2015 Apr;12(4):357–60. doi:10.1038/nmeth.3317 PubMed PMID: 25751142; PubMed Central PMCID: PMC4655817.

28. Liao Y, Smyth GK, Shi W. featureCounts: an efficient general purpose program for assigning sequence reads to genomic features. Bioinformatics. 2014 Apr 1;30(7):923–30. doi:10.1093/bioinformatics/btt656 PubMed PMID: 24227677.

29. Barnett DW, Garrison EK, Quinlan AR, Strömberg MP, Marth GT. BamTools: a C++ API and toolkit for analyzing and managing BAM files. Bioinformatics. 2011 Jun 15;27(12):1691–2. doi:10.1093/bioinformatics/btr174 PubMed PMID: 21493652; PubMed Central PMCID: PMC3106182.

30. Picard Tools [Internet]. Broad Institute; 2019. Available from: http://broadinstitute.github.io/picard/

31. Pedersen BS, Quinlan AR. Who’s Who? Detecting and Resolving Sample Anomalies in Human DNA Sequencing Studies with Peddy. Am J Hum Genet. 2017 Mar 2;100(3):406–13. doi:10.1016/j.ajhg.2017.01.017 PubMed PMID: 28190455; PubMed Central PMCID: PMC5339084.

32. Admati I, Skarbianskis N, Hochgerner H, Ophir O, Yagel S, Solt I, et al. Single-nuclei RNA-sequencing fails to detect molecular dysregulation in the preeclamptic placenta. Placenta. 2025 Jan;159:170–9. doi:10.1016/j.placenta.2024.12.011 PubMed PMID: 39733647.

33. Newman AM, Steen CB, Liu CL, Gentles AJ, Chaudhuri AA, Scherer F, et al. Determining cell type abundance and expression from bulk tissues with digital cytometry. Nat Biotechnol. 2019 Jul;37(7):773–82. doi:10.1038/s41587-019-0114-2 PubMed PMID: 31061481; PubMed Central PMCID: PMC6610714.

34. Horii M, To C, Morey R, Jacobs MB, Li Y, Nelson KK, et al. Histopathologic and Transcriptomic Profiling Identifies Novel Trophoblast Defects in Patients With Preeclampsia and Maternal Vascular Malperfusion. Mod Pathol Off J U S Can Acad Pathol Inc. 2023 Feb;36(2):100035. doi:10.1016/j.modpat.2022.100035 PubMed PMID: 36853788; PubMed Central PMCID: PMC10081686.

35. Ritchie ME, Phipson B, Wu D, Hu Y, Law CW, Shi W, et al. limma powers differential expression analyses for RNA-sequencing and microarray studies. Nucleic Acids Res. 2015 Apr 20;43(7):e47–e47. doi:10.1093/nar/gkv007

36. Robinson MD, McCarthy DJ, Smyth GK. edgeR : a Bioconductor package for differential expression analysis of digital gene expression data. Bioinformatics. 2010 Jan 1;26(1):139–40. doi:10.1093/bioinformatics/btp616

37. Robinson MD, Oshlack A. A scaling normalization method for differential expression analysis of RNA-seq data. Genome Biol. 2010 Mar 2;11(3):R25. doi:10.1186/gb-2010-11-3-r25

38. Law CW, Chen Y, Shi W, Smyth GK. voom: precision weights unlock linear model analysis tools for RNA-seq read counts. Genome Biol. 2014 Feb 3;15(2):R29. doi:10.1186/gb-2014-15-2-r29

39. Xu S, Hu E, Cai Y, Xie Z, Luo X, Zhan L, et al. Using clusterProfiler to characterize multiomics data. Nat Protoc. 2024 Nov;19(11):3292–320. doi:10.1038/s41596-024-01020-z PubMed PMID: 39019974.

40. Yu G, Wang LG, Han Y, He QY. clusterProfiler: an R package for comparing biological themes among gene clusters. Omics J Integr Biol. 2012 May;16(5):284–7. doi:10.1089/omi.2011.0118 PubMed PMID: 22455463; PubMed Central PMCID: PMC3339379.

41. Wu T, Hu E, Xu S, Chen M, Guo P, Dai Z, et al. clusterProfiler 4.0: A universal enrichment tool for interpreting omics data. The Innovation. 2021 Aug;2(3):100141. doi:10.1016/j.xinn.2021.100141

42. Holmgren L, Glaser A, Pfeifer-Ohlsson S, Ohlsson R. Angiogenesis during human extraembryonic development involves the spatiotemporal control of PDGF ligand and receptor gene expression. Development. 1991 Nov;113(3):749–54. doi:10.1242/dev.113.3.749 PubMed PMID: 1668277.

43. Haouzi D, Dechaud H, Assou S, Monzo C, de Vos J, Hamamah S. Transcriptome analysis reveals dialogues between human trophectoderm and endometrial cells during the implantation period. Hum Reprod. 2011 Jun;26(6):1440–9. doi:10.1093/humrep/der075 PubMed PMID: 21427117.

44. Harville EW, Mishra PP, Kähönen M, Raitoharju E, Marttila S, Raitakari O, et al. Reproductive history and blood cell DNA methylation later in life: the Young Finns Study. Clin Epigenetics. 2021 Dec 20;13(1):227. doi:10.1186/s13148-021-01215-1 PubMed PMID: 34930449; PubMed Central PMCID: PMC8690999.

45. Huang X, Li Z, Lei J, Wang D, Zhang Y. Genetic polymorphism in DGCR8 is associated with late onset of preeclampsia. BMC Med Genet. 2019 Sep 4;20(1):151. doi:10.1186/s12881-019-0887-7 PubMed PMID: 31484500; PubMed Central PMCID: PMC6727569.

46. Tsuchida N, Kojima J, Fukuda A, Oda M, Kawasaki T, Ito H, et al. Transcriptomic features of trophoblast lineage cells derived from human induced pluripotent stem cells treated with BMP 4. Placenta. 2020 Jan;89:20–32. doi:10.1016/j.placenta.2019.10.006

47. Workalemahu T, Enquobahrie DA, Gelaye B, Sanchez SE, Garcia PJ, Tekola-Ayele F, et al. Genetic variations and risk of placental abruption: A genome-wide association study and meta-analysis of genome-wide association studies. Placenta. 2018 Jun;66:8–16. doi:10.1016/j.placenta.2018.04.008 PubMed PMID: 29884306; PubMed Central PMCID: PMC5995331.

48. Mousa A. Epigenetic Regulation of Genes Involved in Vascular Dysfunction in Preeclamptic Women [Internet]. Virginia Commonwealth University; 2012. Available from: https://scispace.com/pdf/epigenetic-regulation-of-genes-involved-in-vascular-3wrs1w09ye.pdf

49. Meng SL, Anane-Wae R, Benavente ED, Aherrahrou R. System Genetics Analysis Reveals Sex Differences in Human Aortic Smooth Muscle Gene Expression. Bioinforma Biol Insights. 2024;18:11779322241298592. doi:10.1177/11779322241298592 PubMed PMID: 39619737; PubMed Central PMCID: PMC11605743.

50. Paparini DE, Grasso E, Aguilera F, Arslanian MA, Lella V, Lara B, et al. Sex-specific phenotypical, functional and metabolic profiles of human term placenta macrophages. Biol Sex Differ. 2024 Oct 17;15(1):80. doi:10.1186/s13293-024-00652-w PubMed PMID: 39420346; PubMed Central PMCID: PMC11484421.

51. Muralimanoharan S, Gao X, Weintraub S, Myatt L, Maloyan A. Sexual dimorphism in activation of placental autophagy in obese women with evidence for fetal programming from a placenta-specific mouse model. Autophagy. 2016 May 3;12(5):752–69. doi:10.1080/15548627.2016.1156822

52. Branco MR, King M, Perez-Garcia V, Bogutz AB, Caley M, Fineberg E, et al. Maternal DNA Methylation Regulates Early Trophoblast Development. Dev Cell. 2016 Jan;36(2):152–63. doi:10.1016/j.devcel.2015.12.027

53. Keenen MM, Yang L, Liang H, Farmer VJ, Worota RE, Singh R, et al. Comparative analysis of the syncytiotrophoblast in placenta tissue and trophoblast organoids using snRNA sequencing. eLife. 2025 May 27;13:RP101170. doi:10.7554/eLife.101170 PubMed PMID: 40424181; PubMed Central PMCID: PMC12113261.

54. Xiang Y, Cheng Y, Li X, Li Q, Xu J, Zhang J, et al. Up-regulated expression and aberrant DNA methylation of LEP and SH3PXD2A in pre-eclampsia. PloS One. 2013;8(3):e59753. doi:10.1371/journal.pone.0059753 PubMed PMID: 23544093; PubMed Central PMCID: PMC3609796.

55. Gonzalez TL, Willson BE, Wang ET, Taylor KD, Novoa A, Swarna A, et al. Sexually dimorphic DNA methylation and gene expression patterns in human first trimester placenta. Biol Sex Differ. 2024 Aug 16;15(1):63. doi:10.1186/s13293-024-00629-9 PubMed PMID: 39152463; PubMed Central PMCID: PMC11328442.

56. Couture C, Brien ME, Piché J, Cervantes E, Luu TM, Girard S. Adverse neonatal outcomes are associated with a sex-specific placental inflammatory profile†. Biol Reprod. 2025 Nov 14;113(5):1266–73. doi:10.1093/biolre/ioaf158 PubMed PMID: 40679433; PubMed Central PMCID: PMC12621311.

57. Roland CS, Hu J, Ren CE, Chen H, Li J, Varvoutis MS, et al. Morphological changes of placental syncytium and their implications for the pathogenesis of preeclampsia. Cell Mol Life Sci CMLS. 2016 Jan;73(2):365–76. doi:10.1007/s00018-015-2069-x PubMed PMID: 26496726; PubMed Central PMCID: PMC4846582.

58. Rosario FJ, Powell TL, Gupta MB, Cox L, Jansson T. mTORC1 Transcriptional Regulation of Ribosome Subunits, Protein Synthesis, and Molecular Transport in Primary Human Trophoblast Cells. Front Cell Dev Biol. 2020;8:583801. doi:10.3389/fcell.2020.583801 PubMed PMID: 33324640; PubMed Central PMCID: PMC7726231.

59. Koga K, Mor G. Toll-like receptors at the maternal-fetal interface in normal pregnancy and pregnancy disorders. Am J Reprod Immunol. 2010 Jun;63(6):587–600. doi:10.1111/j.1600-0897.2010.00848.x PubMed PMID: 20367625; PubMed Central PMCID: PMC3025804.

60. Kim YM, Romero R, Oh SY, Kim CJ, Kilburn BA, Armant DR, et al. Toll-like receptor 4: a potential link between “danger signals,” the innate immune system, and preeclampsia? Am J Obstet Gynecol. 2005 Sep;193(3 Pt 2):921–7. doi:10.1016/j.ajog.2005.07.076 PubMed PMID: 16157088.

61. Wang X, Athayde N, Trudinger B. Placental vascular disease and toll-like receptor 4 gene expression. Am J Obstet Gynecol. 2005 Mar;192(3):961–6. doi:10.1016/j.ajog.2004.09.009 PubMed PMID: 15746698.

62. Benouda I, Vaiman D, Miralles F. Trophoblast Fusion in Hypertensive Disorders of Pregnancy and Preeclampsia. Int J Mol Sci. 2025 Mar 21;26(7):2859. doi:10.3390/ijms26072859

63. Li C, Meng Y, Zhou B, Zhang Y, Xia Q, Huang Y, et al. ITGB3 is reduced in pregnancies with preeclampsia and its influence on biological behavior of trophoblast cells. Mol Med. 2024 Dec 25;30(1):275. doi:10.1186/s10020-024-01050-z PubMed PMID: 39721996; PubMed Central PMCID: PMC11670450.

64. He D, Peng X, Xie H, Peng R, Li Q, Guo Y, et al. Genetic Variations in Angiotensinogen Gene and Risk of Preeclampsia: A Pilot Study. J Clin Med. 2023 Feb 14;12(4):1509. doi:10.3390/jcm12041509 PubMed PMID: 36836041; PubMed Central PMCID: PMC9966751.

65. Williams PJ, Broughton Pipkin F. The genetics of pre-eclampsia and other hypertensive disorders of pregnancy. Best Pract Res Clin Obstet Gynaecol. 2011 Aug;25(4):405–17. doi:10.1016/j.bpobgyn.2011.02.007 PubMed PMID: 21429808; PubMed Central PMCID: PMC3145161.

66. Vaughan JE, Walsh SW. Activation of NF-κB in placentas of women with preeclampsia. Hypertens Pregnancy. 2012;31(2):243–51. doi:10.3109/10641955.2011.642436 PubMed PMID: 22380486; PubMed Central PMCID: PMC3542769.

67. Vangrieken P, Al-Nasiry S, Bast A, Leermakers PA, Tulen CBM, Schiffers PMH, et al. Placental Mitochondrial Abnormalities in Preeclampsia. Reprod Sci. 2021 Aug;28(8):2186–99. doi:10.1007/s43032-021-00464-y PubMed PMID: 33523425; PubMed Central PMCID: PMC8289780.

68. Marín R, Chiarello DI, Abad C, Rojas D, Toledo F, Sobrevia L. Oxidative stress and mitochondrial dysfunction in early-onset and late-onset preeclampsia. Biochim Biophys Acta BBA -Mol Basis Dis. 2020 Dec;1866(12):165961. doi:10.1016/j.bbadis.2020.165961

69. Matsubara S, Minakami H, Sato I, Saito T. Decrease in cytochrome c oxidase activity detected cytochemically in the placental trophoblast of patients with pre-eclampsia. Placenta. 1997 May;18(4):255–9. doi:10.1016/S0143-4004(97)80059-8

70. Shi Z, Long W, Zhao C, Guo X, Shen R, Ding H. Comparative proteomics analysis suggests that placental mitochondria are involved in the development of pre-eclampsia. PloS One. 2013;8(5):e64351. doi:10.1371/journal.pone.0064351 PubMed PMID: 23671712; PubMed Central PMCID: PMC3650054.

71. Voros C, Stavros S, Sapantzoglou I, Mavrogianni D, Daskalaki MA, Theodora M, et al. The Role of Placental Mitochondrial Dysfunction in Adverse Perinatal Outcomes: A Systematic Review. J Clin Med. 2025 May 29;14(11):3838. doi:10.3390/jcm14113838 PubMed PMID: 40507600; PubMed Central PMCID: PMC12155783.

72. Hu XQ, Zhang L. Mitochondrial Dysfunction in the Pathogenesis of Preeclampsia. Curr Hypertens Rep. 2022 Jun;24(6):157–72. doi:10.1007/s11906-022-01184-7 PubMed PMID: 35254588; PubMed Central PMCID: PMC10054606.

73. Kelly A, Chan J, Powell TL, Cox LA, Jansson T, Rosario FJ. Maternal obesity alters the placental transcriptome in a fetal sex-dependent manner. Front Cell Dev Biol. 2023 Jun 15;11:1178533. doi:10.3389/fcell.2023.1178533

74. Salazar-Petres E, Pereira-Carvalho D, Lopez-Tello J, Sferruzzi-Perri AN. Maternal and Intrauterine Influences on Feto-Placental Growth Are Accompanied by Sexually Dimorphic Changes in Placental Mitochondrial Respiration, and Metabolic Signalling Pathways. Cells. 2023 Mar 3;12(5):797. doi:10.3390/cells12050797

75. Meakin AS, Cuffe JSM, Darby JRT, Morrison JL, Clifton VL. Let’s Talk about Placental Sex, Baby: Understanding Mechanisms That Drive Female-and Male-Specific Fetal Growth and Developmental Outcomes. Int J Mol Sci. 2021 Jun 15;22(12):6386. doi:10.3390/ijms22126386 PubMed PMID: 34203717; PubMed Central PMCID: PMC8232290.

76. Choi SJ, Oh S young, Kim JH, Sadovsky Y, Roh CR. Increased expression of N-myc downstream-regulated gene 1 (NDRG1) in placentas from pregnancies complicated by intrauterine growth restriction or preeclampsia. Am J Obstet Gynecol. 2007 Jan;196(1):45.e1-7. doi:10.1016/j.ajog.2006.08.029 PubMed PMID: 17240229.

77. Fu Y, Wei J, Dai X, Ye Y. Increased NDRG1 expression attenuate trophoblast invasion through ERK/MMP-9 pathway in preeclampsia. Placenta. 2017 Mar;51:76–81. doi:10.1016/j.placenta.2017.01.126 PubMed PMID: 28292472.

78. Zhang Y, Chu J, Yu Z, Gong Y. Single cell RNA-sequencing reveals BHLHE40 and NDRG1 as key regulatory molecules modulating the trophoblastic senescence in preeclampsia. Placenta. 2025 Aug;168:74–87. doi:10.1016/j.placenta.2025.06.009 PubMed PMID: 40517485.

79. Wong GP, Hartmann S, Simmons DG, Ellis S, Nonn O, Cannon P, et al. Trophoblast Side-Population Markers are Dysregulated in Preeclampsia and Fetal Growth Restriction. Stem Cell Rev Rep. 2024 Oct;20(7):1954–70. doi:10.1007/s12015-024-10764-w PubMed PMID: 39028417; PubMed Central PMCID: PMC11445292.

80. Wang C, Ju H, Zhou L, Zhu Y, Wu L, Deng X, et al. TET3-mediated novel regulatory mechanism affecting trophoblast invasion and migration: Implications for preeclampsia development. Placenta. 2024 Mar 6;147:31–41. doi:10.1016/j.placenta.2024.01.010 PubMed PMID: 38295560.

81. Gonzalez TL, Sun T, Koeppel AF, Lee B, Wang ET, Farber CR, et al. Sex differences in the late first trimester human placenta transcriptome. Biol Sex Differ. 2018 Jan 15;9(1):4. doi:10.1186/s13293-018-0165-y PubMed PMID: 29335024; PubMed Central PMCID: PMC5769539.

82. Zhu H, Peng B, Klausen C, Yi Y, Li Y, Xiong S, et al. NPFF increases fusogenic proteins syncytin 1 and syncytin 2 via GCM1 in first trimester primary human cytotrophoblast cells. FASEB J Off Publ Fed Am Soc Exp Biol. 2020 Jul;34(7):9419–32. doi:10.1096/fj.201902978R PubMed PMID: 32501590.

83. Arroyo J, Price M, Straszewski-Chavez S, Torry RJ, Mor G, Torry DS. XIAP protein is induced by placenta growth factor (PLGF) and decreased during preeclampsia in trophoblast cells. Syst Biol Reprod Med. 2014 Oct;60(5):263–73. doi:10.3109/19396368.2014.927540 PubMed PMID: 25003840.

84. Zheng S, Chen X, Tang W, Yang D, Han Q. Identification and Validation of DLD as a Cuproptosis-Associated Biomarker in Preeclampsia. FASEB J Off Publ Fed Am Soc Exp Biol. 2026 Apr 30;40(8):e71787. doi:10.1096/fj.202504941R PubMed PMID: 41996226; PubMed Central PMCID: PMC13089668.

85. Plaisier SB, Taschereau R, Wong JA, Graeber TG. Rank-rank hypergeometric overlap: identification of statistically significant overlap between gene-expression signatures. Nucleic Acids Res. 2010 Sep;38(17):e169. doi:10.1093/nar/gkq636 PubMed PMID: 20660011; PubMed Central PMCID: PMC2943622.

86. Dackor J, Strunk KE, Wehmeyer MM, Threadgill DW. Altered trophoblast proliferation is insufficient to account for placental dysfunction in Egfr null embryos. Placenta. 2007;28(11–12):1211–8. doi:10.1016/j.placenta.2007.07.005 PubMed PMID: 17822758; PubMed Central PMCID: PMC2121666.

87. Zhou Y, Fisher SJ, Janatpour M, Genbacev O, Dejana E, Wheelock M, et al. Human cytotrophoblasts adopt a vascular phenotype as they differentiate. A strategy for successful endovascular invasion? J Clin Invest. 1997 May 1;99(9):2139–51. doi:10.1172/JCI119387 PubMed PMID: 9151786; PubMed Central PMCID: PMC508044.

88. Zhou Y, Damsky CH, Fisher SJ. Preeclampsia is associated with failure of human cytotrophoblasts to mimic a vascular adhesion phenotype. One cause of defective endovascular invasion in this syndrome? J Clin Invest. 1997 May 1;99(9):2152–64. doi:10.1172/JCI119388 PubMed PMID: 9151787; PubMed Central PMCID: PMC508045.

89. Gao L, Mathur V, Tam SKM, Zhou X, Cheung MF, Chan LY, et al. Single-cell analysis reveals transcriptomic and epigenomic impacts on the maternal-fetal interface following SARS-CoV-2 infection. Nat Cell Biol. 2023 Jul;25(7):1047–60. doi:10.1038/s41556-023-01169-x PubMed PMID: 37400500; PubMed Central PMCID: PMC10344786.

90. Oh JH, Rizzuto G, Elkin R, Weistuch C, Norton L, Dveksler G, et al. Pregnancy-specific glycoproteins as potential drug targets for female lung adenocarcinoma patients. Brief Funct Genomics. 2025 Jan 15;24:elaf004. doi:10.1093/bfgp/elaf004

91. Chang CL, Chang CY, Lee DX, Cheng PJ. Characterization of Human Pregnancy Specific Glycoprotein (PSG) Gene Copy Number Variations in Pre-eclampsia Patients. Adv Exp Med Biol. 2016;924:63–5. doi:10.1007/978-3-319-42044-8_12 PubMed PMID: 27753020.

92. Zhou Y, Damsky CH, Chiu K, Roberts JM, Fisher SJ. Preeclampsia is associated with abnormal expression of adhesion molecules by invasive cytotrophoblasts. J Clin Invest. 1993 Mar;91(3):950–60. doi:10.1172/JCI116316 PubMed PMID: 7680671; PubMed Central PMCID: PMC288047.

93. Zhang B, Song C, Zhou B, Zhang J, Dong W, Zhang Y, et al. CTNNB1 and CDH1 Regulate Trophoblast Cell Adhesion and Junction Formation in Yak Placental Tissue at Different Gestational Stages. Anim Open Access J MDPI. 2025 Mar 19;15(6):876. doi:10.3390/ani15060876 PubMed PMID: 40150405; PubMed Central PMCID: PMC11939409.

94. Du L, Kuang L, He F, Tang W, Sun W, Chen D. Mesenchymal-to-epithelial transition in the placental tissues of patients with preeclampsia. Hypertens Res Off J Jpn Soc Hypertens. 2017 Jan;40(1):67–72. doi:10.1038/hr.2016.97 PubMed PMID: 27511055.

95. Kurioka D, Takeshita F, Tsuta K, Sakamoto H, Watanabe S ichi, Matsumoto K, et al. NEK9-dependent proliferation of cancer cells lacking functional p53. Sci Rep. 2014 Aug 18;4:6111. doi:10.1038/srep06111 PubMed PMID: 25131192; PubMed Central PMCID: PMC4135330.

96. Tompkins VS, Hagen J, Frazier AA, Lushnikova T, Fitzgerald MP, Di Tommaso A, et al. A Novel Nuclear Interactor of ARF and MDM2 (NIAM) That Maintains Chromosomal Stability. J Biol Chem. 2007 Jan;282(2):1322–33. doi:10.1074/jbc.M609612200

97. Oksanen M, Mastropasqua F, Mazan-Mamczarz K, Martindale JL, Ye X, Arora A, et al. Molecular interactome of HNRNPU reveals regulatory networks in neuronal differentiation and DNA methylation. Nucleic Acids Res. 2026 Feb 5;54(4):gkag107. doi:10.1093/nar/gkag107

98. Illsley NP, Zamudio S. Epithelial-mesenchymal plasticity in trophoblast differentiation. Placenta. 2026 Oct;184:74–85. doi:10.1016/j.placenta.2026.02.009 PubMed PMID: 41741289.

99. Sood R, Kalloway S, Mast AE, Hillard CJ, Weiler H. Fetomaternal cross talk in the placental vascular bed: control of coagulation by trophoblast cells. Blood. 2006 Apr 15;107(8):3173–80. doi:10.1182/blood-2005-10-4111 PubMed PMID: 16380449; PubMed Central PMCID: PMC1895751.

100. Rusidzé M, Gargaros A, Fébrissy C, Dubucs C, Weyl A, Ousselin J, et al. Estrogen Actions in Placental Vascular Morphogenesis and Spiral Artery Remodeling: A Comparative View between Humans and Mice. Cells. 2023 Feb 14;12(4):620. doi:10.3390/cells12040620 PubMed PMID: 36831287; PubMed Central PMCID: PMC9954071.

101. Huppertz B, Kadyrov M, Kingdom JCP. Apoptosis and its role in the trophoblast. Am J Obstet Gynecol. 2006 Jul;195(1):29–39. doi:10.1016/j.ajog.2005.07.039 PubMed PMID: 16579915.

102. Allaire AD, Ballenger KA, Wells SR, McMahon MJ, Lessey BA. Placental apoptosis in preeclampsia. Obstet Gynecol. 2000 Aug;96(2):271–6. doi:10.1016/s0029-7844(00)00895-4 PubMed PMID: 10908776.

103. Leung DN, Smith SC, To KF, Sahota DS, Baker PN. Increased placental apoptosis in pregnancies complicated by preeclampsia. Am J Obstet Gynecol. 2001 May;184(6):1249–50. doi:10.1067/mob.2001.112906 PubMed PMID: 11349196.

104. Ishihara N, Matsuo H, Murakoshi H, Laoag-Fernandez JB, Samoto T, Maruo T. Increased apoptosis in the syncytiotrophoblast in human term placentas complicated by either preeclampsia or intrauterine growth retardation. Am J Obstet Gynecol. 2002 Jan;186(1):158–66. doi:10.1067/mob.2002.119176 PubMed PMID: 11810103.

105. Vangrieken P, Al-Nasiry S, Bast A, Leermakers PA, Tulen CBM, Janssen GMJ, et al. Hypoxia-induced mitochondrial abnormalities in cells of the placenta. PloS One. 2021;16(1):e0245155. doi:10.1371/journal.pone.0245155 PubMed PMID: 33434211; PubMed Central PMCID: PMC7802931.

106. Liberzon A, Birger C, Thorvaldsdóttir H, Ghandi M, Mesirov JP, Tamayo P. The Molecular Signatures Database Hallmark Gene Set Collection. Cell Syst. 2015 Dec;1(6):417–25. doi:10.1016/j.cels.2015.12.004

107. Kleinrouweler CE, Van Uitert M, Moerland PD, Ris-Stalpers C, Van Der Post JAM, Afink GB. Differentially Expressed Genes in the Pre-Eclamptic Placenta: A Systematic Review and Meta-Analysis. Oudejans C, editor. PLoS ONE. 2013 Jul 12;8(7):e68991. doi:10.1371/journal.pone.0068991

108. Zeisler H, Llurba E, Chantraine F, Vatish M, Staff AC, Sennström M, et al. Predictive Value of the sFlt-1:PlGF Ratio in Women with Suspected Preeclampsia. N Engl J Med. 2016 Jan 7;374(1):13–22. doi:10.1056/NEJMoa1414838

109. Middendorf L, Gellhaus A, Iannaccone A, Köninger A, Dathe AK, Bendix I, et al. The Impact of Increased Maternal sFlt-1/PlGF Ratio on Motor Outcome of Preterm Infants. Front Endocrinol. 2022 Jun 30;13:913514. doi:10.3389/fendo.2022.913514

110. Vogtmann R, Bao M, Dewan MV, Riedel A, Kimmig R, Felderhoff-Müser U, et al. Growth-Restricted Fetuses and Offspring Reveal Adverse Sex-Specific Metabolic Responses in Preeclamptic Mice Expressing Human sFLT1. Int J Mol Sci. 2023 Apr 7;24(8):6885. doi:10.3390/ijms24086885 PubMed PMID: 37108049; PubMed Central PMCID: PMC10139224.

111. Andersen LB, Jørgensen JS, Herse F, Andersen MS, Christesen HT, Dechend R. The association between angiogenic markers and fetal sex: Implications for preeclampsia research. J Reprod Immunol. 2016 Sep;117:24–9. doi:10.1016/j.jri.2016.05.005

112. Taylor BD, Haggerty CL, Ness RB, Hougaard DM, Skogstrand K, Roberts JM, et al. Fetal sexual dimorphism in systemic soluble fms-like tyrosine kinase 1 among normotensive and preeclamptic women. Am J Reprod Immunol. 2018 Nov;80(5):e13034. doi:10.1111/aji.13034 PubMed PMID: 30106204.

113. Jennewein DM, Lee J, Kurtz C, Dizon W, Shaeffer I, Chapman A, et al. The Sol Supercomputer at Arizona State University. In: Practice and Experience in Advanced Research Computing [Internet]. Portland OR USA: ACM; 2023 [cited 2026 Nov 4]. p. 296–301. Available from: https://dl.acm.org/doi/10.1145/3569951.3597573 doi:10.1145/3569951.3597573

